# Involvement of RNase J in CRISPR RNA maturation in the cyanobacterium *Synechocystis* sp. PCC 6803

**DOI:** 10.1101/2025.02.21.638496

**Authors:** Raphael Bilger, Friedel Drepper, Bettina Knapp, Tanja Berndt, Helena Landerer, Harald Putzer, Pitter Huesgen, Wolfgang R. Hess

**Affiliations:** University of Freiburg, Faculty of Biology, Genetics and Experimental Bioinformatics, Schänzlestr. 1, D-79104 Freiburg, Germany; University of Freiburg, Faculty of Biology, Biochemistry and Functional Proteomics, Schänzlestr. 1, D-79104 Freiburg, Germany; Expression Génétique Microbienne (EGM), CNRS, Université Paris Cité, Institut de Biologie Physico-Chimique, F-75005 Paris, France; University of Freiburg, CIBSS Centre for Integrative Biological Signalling Studies, D-79104 Freiburg, Germany

**Keywords:** CRISPR-Cas systems, cyanobacteria, degradosome, gene expression regulation, ribonuclease J, RNA structure, RNA:RNA interaction

## Abstract

Many bacteria and archaea use CRISPR-Cas systems, which provide RNA-based, adaptive, and inheritable immune defenses against invading viruses and other foreign genetic elements. The proper processing of CRISPR guide RNAs (crRNAs) is a crucial step in the maturation of the defense complexes and is frequently performed by specialized ribonucleases encoded by *cas* genes. However, some systems employ enzymes associated with degradosome or housekeeping functions, such as RNase III or the endoribonuclease RNase E. Here, the endo- and 5′-exoribonuclease RNase J was identified as an additional enzyme involved in crRNA maturation, acting jointly with RNase E in the crRNA maturation of a type III-Bv CRISPR-Cas system, and possibly together with a further RNase in the cyanobacterium *Synechocystis* sp. PCC 6803. Co-IP experiments revealed a small set of proteins that were co-enriched with RNase J, among them the exoribonuclease polyribonucleotide nucleotidyltransferase (PNPase). Despite a measured, strong 3’ exonucleolytic activity of the recombinant enzyme, PNPase was not confirmed to contribute to crRNA maturation. However, the co-IP results indicate that PNPase in *Synechocystis* is an enzyme that can recruit either RNase E or RNase J, together with additional proteins.

## Introduction

Bacteria are exposed to many environmental changes that require rapid acclimation responses, including the regulation of gene expression. One mechanism is the post-transcriptional regulation of mRNA decay involving the activity of one of the major ribonucleases (RNases), RNase E/G, RNase Y, and/or RNase J. In all bacteria, at least one homolog of these RNases can be found (Laalami, Zig and Putzer 2014; Aït-Bara and Carpousis 2015).

RNases are often the scaffold for an mRNA degradation machinery, the degradosome. In *E. coli*, RNase E possesses binding domains in its C-terminal region to recruit the exoribonuclease polyribonucleotide nucleotidyltransferase (PNPase), the RNA helicase RhlB and the glycolytic enzyme enolase (Vanzo *et al*. 1998). In the Gram-positive bacterium *B. subtilis*, the degradosome complex is assembled around RNase Y, which, similarly to the *E.coli* degradosome, interacts with PNPase, the helicase CshA and enzymes involved in glycolysis (Commichau *et al*. 2009; Lehnik-Habrink *et al*. 2010; Newman *et al*. 2012).

An RNase E-based RNA degradosome exists also in cyanobacteria (Zhang, Hess and Zhang 2022). It was first shown in the filamentous cyanobacterium *Nostoc* (*Anabaena*) sp. PCC 7120 that the noncatalytic region of RNase E recruited PNPase, and that the two proteins interact through a widely conserved nonapeptide within the RNase E C-terminal region (subregion C4) (Zhang *et al*. 2014). Later, the 3’ exoribonuclease RNase II was found to be another RNase E-associated ribonuclease in this organism (Zhou *et al*. 2020), and there is strong evidence for further additional interacting proteins, such as the RNA helicase CrhB, enolase (Yan *et al*. 2020), and the conserved protein RebA (Liu *et al*. 2024). RebA is an inhibitor of RNase E catalytic activity and is universally conserved in cyanobacteria (Liu *et al*. 2024).

Besides mRNA degradation, the degradosome enzymes play important roles in other biological processes, such as in the control of translation (Redko *et al*. 2013) or the maturation of CRISPR-related transcripts (crRNAs) (Behler *et al*. 2018). The type III-A CRISPR/Cas system of *Staphylococcus epidermidis* relies on the degradosome associated PNPase and RNase J2 to promote crRNA maturation and for efficient clearance of phage-derived nucleic acids (Chou-Zheng and Hatoum-Aslan 2019).

In the cyanobacterial model *Synechocystis* sp. PCC 6803 (hereafter *Synechocystis*), three distinct CRISPR/Cas systems were described. These are a type I-D (CRISPR1), a type III-Dv (CRISPR2), and a III-Bv (CRISPR3) system, which are highly expressed under a variety of conditions and active in interference assays (Scholz *et al*. 2013, 2019; Kieper *et al*. 2018; McBride *et al*. 2020; Bilger *et al*. 2024; Schwartz *et al*. 2024). While the CRISPR1 and CRISPR2 systems depend on Cas6 endoribonucleases for the maturation of crRNAs (Scholz *et al*. 2013; Reimann *et al*. 2017), no such enzyme is affiliated with the CRISPR3 system. Instead, the CRISPR3 system depends on the major ribonuclease RNase E to process crRNAs in this system. It catalyzes a single cleavage within the repeats, directly generating the 13 and 14 nt long 5’ repeat handles (Behler *et al*. 2018). However, the sequences between the spacer ends and the 5’ handle of the next following crRNA disappear and also do not accumulate as individual fragments. These observations and the accumulation of mature crRNAs exclusively 48 and 42 nt in length suggested the involvement of at least one additional ribonuclease in the maturation of CRISPR3-derived crRNAs (Scholz *et al*. 2013; Behler *et al*. 2018).

In addition to RNase E (*slr1129*), RNase J (*slr0551*) is present in *Synechocystis*, a combination that is conserved in cyanobacteria (Laalami and Putzer 2011). RNase E has been analyzed in more depth than RNase J, but it is clear that both enzymes target a wide range of transcripts (Cavaiuolo *et al*. 2020; Hoffmann *et al*. 2021, 2024). Neither the gene encoding RNase E nor the gene for RNase J can be deleted in *Synechocystis* indicating their essentiality. However, *Synechocystis* contains multiple chromosome copies (Zerulla, Ludt and Soppa 2016), making partial gene deletions possible to generate knockdown mutants with a reduced gene copy number. Analysis of such a partial *rnj* deletion (strain δ*rnj*) indicated that the maturation of crRNAs from the CRISPR3 system could involve RNase J (Cavaiuolo *et al*. 2020).

RNase J was first discovered in *Bacillus subtilis* (Even *et al*. 2005), but is wide-spread also in other bacteria, in many archaea (Hasenöhrl, Konrat and Bläsi 2011; Levy *et al*. 2011; Zheng *et al*. 2017), and also present in plant and algal chloroplasts (Chen, Zou and Zhao 2015; Liponska *et al*. 2018; Hotto, Stern and Schuster 2020). RNase J belongs to the β-CASP protein family within the superfamily of metallo-β-lactamase fold proteins (Callebaut *et al*. 2002), with two beta lactamase domains separated by a β-CASP domain (Condon and Gilet 2011). The enzyme possesses an endonuclease and a unique 5’-3’ exonuclease activity (Mathy *et al*. 2007), the latter previously thought to be absent in prokaryotes (Alifano, Bruni and Carlomagno 1994; Deana and Belasco 2005). Despite these insights, RNase J is still less well investigated in comparison to RNase E (Ehretsmann, Carpousis and Krisch 1992; Mackie 2013; Zhang *et al*. 2014; Chao *et al*. 2017; Hoffmann *et al*. 2021, 2024; Liu *et al*. 2024), or RNase Y (Shahbabian *et al*. 2009; Lehnik-Habrink *et al*. 2011; Marincola *et al*. 2012; Scornet *et al*. 2024).

Here, we investigated if RNase J is involved in the maturation of pre-crRNA in *Synechocystis* and identified putative cleavage sites. Through co-immunoprecipitation (co-IP) coupled with mass spectrometry we identified RNase J interaction partners, pointing at possible further proteins interacting with RNase J in *Synechocystis*.

## Methods

### Strains and cultivation

*Synechocystis* cultures were grown at 30°C in 50 mL copper-depleted BG11 medium (Rippka *et al*. 1979) supplemented with 20 mM TES (N-[Tris-(hydroxymethyl)-methyl]-2-aminoethane sulfonic acid) under continuous illumination with white light at 30 – 50 μmol photons m^-2^ s^-1^ and constant shaking. The cultures were supplied with appropriate antibiotics, namely kanamycin (50 µg/mL) and gentamicin (2 µg/mL) for *Synechocystis* strains carrying pVZ322 (Zinchenko *et al*. 1999) constructs, and kanamycin (100 µg/mL) and spectinomycin (200 µg/mL) for the strains carrying the spectinomycin resistance cassette and a pVZ322 vector. For co-IP assays, four biological replicates of strains overexpressing N-or C-terminally tagged RNase J and GFP controls were cultivated in a high-density cultivation system (HD100 Cultivator from CellDEG GmbH) in 100 mL of fresh water organism (FWO) medium (Migur *et al*. 2021) at a starting OD_750_ of 0.9 – 1.0. FWO medium consists of 50 mM NaNO_3_, 15 mM KNO_3_, 2 mM MgSO_4_, 0.5 mM CaCl_2_, 0.025 mM H_3_BO_3_, 0.15 mM FeCl_3_/Na_2_EDTA, 1.6 mM KH_2_PO_4_, 2.4 mM K_2_HPO_4_, 10 mM NaHCO_3_, 10 μM MnCl_2_, 1 μM ZnSO_4_, 0.1 μM Na_2_MoO_4_, 0.01 μM CuSO_4_ and 0.03 μM CoCl_2_. High CO_2_ concentration were delivered from a carbonate buffer (8.29 g K_2_CO_3_ and 54.06 g KHCO_3_ in 200 mL deionized water) according to manufactures instructions. Kanamycin and gentamicin were added as described (Migur *et al*. 2021). RNase J or GFP expression was induced with 2 µM of Cu^2+^ or 6 mg/mL of L-rhamnose, according to the expression system, for 24 h before harvesting (final OD_750_ of approximately 25 – 30). An equivalent of 1000 OD of cells were collected by centrifugation (3,200 × g, 10 min, at room temperature). Pellets were stored at -70°C until further processing.

### Construction of RNase J expression vectors

We constructed two different systems for the expression of 3×FLAG-tagged RNase J. First, we constructed the pVZ322-P*_petE_*-*slr0551* vector with and without a C-terminal 3×FLAG tag. The *slr0551* native gene and the *petE* gene promoters were amplified from *Synechocystis* genomic DNA using primer pairs RB1/ RB2 and RB3/RB4, respectively. The PCR products were separated on 0.8% agarose gels, visualized with ethidium bromide and correctly-sized fragments were excised and purified with the NucleoSpin Gel and PCR Clean-up kit (Machery-Nagel GmbH), according to manufacturer’s instructions. Then, the fragments were ligated by AQUA cloning (Beyer *et al*. 2015) into pUC19 reverse-amplified by PCR with primers RB5/ RB6 and transformed into chemically competent *E. coli* DH5α cells (NEB). After confirming correct insert sequences, P*_petE_*-*slr0551* was amplified using primers RB7/ RB8, or primers RB9/ RB10 to generate fragments for the inducible RNase J expression with and without a C-terminally 3×FLAG tag. The fragments were ligated into *Pst*I/*Xba*I-digested pVZ322 via AQUA cloning. The final pVZ322 constructs were introduced into the *Synechocystis* wild type, WT-Spc and δ*rnj* (Cavaiuolo *et al*. 2020) strains by tri-parental mating (Scholz *et al*. 2013). Constructs were verified via PCR and sequencing.

To construct plasmid pVZ322-P*_rha_*-3×FLAG-*slr0551* for the expression of RNase J with an N-terminal 3×FLAG tag under the control of an rhamnose-inducible promoter (Kelly *et al*. 2018, 2019), primers RB11/ RB12 were used to amplify a 3×*flag*-*slr0551* fragment. In parallel, the backbone was reverse-amplified from plasmid pUC19P_rha_YFP3×FLAG (Hemm *et al*. 2024) using primer pair RB13/ RB14 . After gel separation and excision, the two amplicons were treated with FastDigest *Dpn*I (Thermo Fisher Scientific) to eliminate template DNA. The construct was assembled via AQUA cloning (Beyer *et al*. 2015) by transformation into DH5α chemical competent *E. coli* cells (NEB). Correct insertion of the *slr0551* fragment was checked by colony PCR with primer pair RB15/ RB16 using One*Taq* polymerase (NEB) and positive clones were sequenced by IDT (Integrated DNA Technologies, Inc). Correct constructs were selected for PCR with primer pair RB15/ RB16 using PCRBio HiFi polymerase (PCR Biosystems Ltd.) and cloned into *Xmn*I-digested pVZ322s (pVZ322 lacking the kanamycin cassette (Kaltenbrunner *et al*. 2023)) by AQUA cloning. The verified construct was then introduced in *Synechocystis* wild type via tri-parental mating (Scholz *et al*. 2013). Correct clones were identified by PCR and verified by sequencing. Primers used in this work are listed in **Supplementary Table S1**.

### RNA isolation

*Synechocystis* wild type, the strain with spectinomycin cassette integrated into a neutral site (in the *ssl0452-slr0271* intergenic region, strain WT-Spc (Cavaiuolo *et al*. 2020)), and δ*rnj* strains with and without pVZ322 constructs for RNase J expression were cultivated in copper depleted BG11 with appropriate antibiotics. When the cultures reached an OD_750_ of approximately 0.5, they were induced with 2 µM CuSO_4_ and collected by vacuum filtration on hydrophilic polyethersulfone filters (Pall Supor®-800, 0.8 μm) at time point 0 (before induction), and 24 and 48 h thereafter. The filter was transferred to a tube containing 1 mL of PGTX (39.6 g phenol, 6.9 g glycerol, 0.1 g 8-hydroxyquinoline, 0.58 g EDTA, 0.8 g sodium acetate, 9.5 g guanidine thiocyanate, 4.6 g guanidine hydrochloride) (Pinto *et al*. 2009), snap frozen in liquid nitrogen and stored at - 80°C. For the RNA extraction, the samples were incubated for 15 min at 65°C, 1 mL of chloroform/isoamyl alcohol (IAA) (24:1) was added and the samples were incubated for 10 min at room temperature. Phase separation was performed by centrifugation for 5 min at 3,270 × g at room temperature. The upper aqueous phase was transferred to a new tube, mixed with one volume of chloroform/IAA (24:1) and centrifuged again. The aqueous phase was transferred to a new tube, mixed with one volume of isopropanol and precipitated overnight at -20°C. The precipitated RNA was pelleted by centrifugation (13,000 × g, 30 min, 4°C). The pellet was washed with 700 μL of 70% ethanol (13,000 × g, 10 min, 4°C), air-dried and resuspended in 20 – 40 μL of nuclease-free water. RNA concentration was determined using a spectrophotometer (NanoDrop ND-1000, Peqlab). The RNA quality was assessed by loading 10 µg of RNA on an 8 M urea, 10% polyacrylamide (PAA)-Tris-Borate-EDTA (TBE) gel.

### Northern analysis

Northern hybridization and analysis was performed as described previously (Bilger *et al*. 2024). Primers RB17 and RB18 were used to generate the template for probe generation used in the hybridization of CRISPR3 crRNA, covering the region from spacer 1 to spacer 4. Desoxyoligonucleotide RB19 was used for quantification of the 5S RNA.

### Primer extension assay

Primer extension was performed as described previously (Behler *et al*. 2018) using the primers RB20 and RB21 (called PrimerExt_Ladder_fwd and PrimerExt_S3_rev in Behler *et al*. (2018)) to target the same region in the CRISPR3 pre-crRNA transcript, and using RNA from *Synechocystis* WT-Spc or δ*rnj* strains, carrying different pVZ322 constructs. The cells were harvest 24 h post-induction and RNA was isolated as described above.

### RNase J co-IP and proteomic sample preparation

Cell pellets from CellDEG cultures were resuspended in 1.5 mL FLAG buffer (50 mM Hepes-NaOH pH 7.5 mM MgCl_2_, 25 mM CaCl_2_, 150 mM NaCL, 10% glycerol, 0.1% Tween-20 and cOmplete Protease inhibitor (Roche)) and split in three screw-cap tubes containing 250 µL glass beads (Ø 0.1-0.25 mm, Retsch). The cells were lysed using a prechilled Precellys homogenizer (Bertin Technologies). To remove cell debris and glass beads, the samples were centrifuged at 13,000 × g for 30 min at 4°C. The supernatant (soluble crude extract) was incubated with 100 µL of packed anti-FLAG^®^ M2 magnetic beads (Sigma Aldrich), for 2 hours at 4°C under gentle rotation. The beads were washed 6 times with 2 mL of FLAG-buffer using DynaMag™-2 magnetic rack (Thermo Fisher). The attached proteins were eluted with 250 µL TBS containing 0.4 mg/mL FLAG peptide as competitor. The samples were incubated for 30 min at 4°C under gentle rotation. The supernatant was taken as elution fraction.

For MS analysis, 10 µg of protein per sample was prepared by adding 1% SDS and 5 mM DTT, followed by an incubation at 60°C for 30 min with shaking at 1,000 rpm. in a ThermoMixer^®^ (Eppendorf). Reduced disulfides were alkylated by adding 0.1 vol of 2-chloroacetamide (200 mM) and incubation for 15 min at 37°C and shaking. The alkylation reaction was quenched by adding 5 mM of DTT. The protein clean-up and digestion was performed following the SP3 protocol (Hughes *et al*. 2019). To each sample, 100 µg of a 1:1 mixture of Sera-Mag SpeedBeads suspension (prepared to a 50 µg/µL, GE Healthcare^®^, cat. No. 45152105050250 and 65152105050250) was added. To induce binding of the proteins to the beads, 4 vol of 100% ethanol were added and incubated in a ThermoMixer^®^ at 24°C for 5 min at 1,000 rpm. Tubes were placed in a magnetic rack and the beads were washed 3 times with 180 µL of 80% ethanol. Beads were resuspended in 75 µl of 100 mM ammonium bicarbonate, pH 8 containing 0.24 µg of sequencing grade, modified trypsin (Promega), sonicated for 30 s in a water bath to disaggregate the beads and incubated for 18 h at 37°C in a ThermoMixer^®^ at 1,000 rpm. After digestion, the samples were centrifuged at 20,000 × g for 1 min at RT. The tubes were placed in the magnetic rack and the supernatant, containing the digested proteins, was transferred to a new tube. Samples were acidified by adding formic acid to a final concentration of approximately 2% to obtain a pH < 3. Next, AttractSPE Tips RPS T1 200 µL (Affinisep) were equilibrated with 50 µL methanol centrifuged for 2 min at 1,000 × g, followed by the addition of 50 µL 0.5% acetic acid and another centrifugation. The acidified samples were loaded onto the equilibrated tips and centrifuged. Samples were washed with 30 µL and 60 µL of 0.1% formic acid and 80% acetonitrile. Samples were eluted with 65 µL of freshly prepared 5% ammonium hydroxide and 60% acetonitrile in water. Peptides were eluted by centrifugation for 3 min at 500 × g into glass vials. The elution was dried in SpeedVac set at 30°C and stored at - 20°C.

### Liquid chromatography and electrospray MS

For LC-MS analysis, an UltiMate^TM^ 3000 RSLCnano system was online coupled to an Exploris 480 mass spectrometer (both Thermo Fisher Scientific, Dreieich, Germany). The peptide mixture was washed and preconcentrated on a μPAC™ C18 trapping column (PharmaFluidics) with a flow rate of 10 µL/min and peptides were separated on a μPAC™ C18 pillar array column (50 cm bed length, PharmaFluidics) using a binary buffer system consisting of A (0.1% formic acid) and B (86% acetonitrile, 0.1% formic acid). With a flow rate of 0.5 µL/min peptides were eluted applying an isocratic gradient from 1% to 4% B in 2 min, 4% to 25% B in 24 min, 25% to 44% B in 12 min, 44% to 90% in 2 min, and 90% B for 4 min. For electrospray ionization of peptides, a Nanospray Flex ion source (Thermo Fisher Scientific) with a liquid junction PST-HV-NFU (MS Wil, The Netherlands) and a fused silica emitter (20 µm inner diameter, 360 µm outer diameter, MicroOmics Technologies LLC) with a source voltage of +1,800 V and an ion transfer tube temperature of 275°C was used. Cycles of data-independent acquisition (DIA) consisted of: one overview spectrum (RF lens of 40%, normalized AGC target of 300%, maximum injection time of 45 ms, *m/z* range of 350 to 1,400, resolution of 120,000) recorded in profile mode followed by MS2 fragment spectra (RF lens of 50%, normalized AGC target of 1000%, maximum injection time of 54 ms) generated sequentially by higher-energy collision-induced dissociation (HCD) at a normalized energy of 28% in a precursor *m/z* range from 361 to 450 in 6 windows of 14 *m/z* isolation width, in a precursor *m/z* range from 450 to 800 in 50 windows of 7 *m/z* isolation width and in a precursor *m/z* range from 800 to 1,100 in 21 windows of 14 *m/z* isolation width, overlapping by 1 *m/z* and recorded at a resolution of 30,000 in profile mode.

### MS data analysis

MS raw data were converted to mzML format using the ProteoWizard software, version 3.0.21229 (Chambers *et al*. 2012). For protein and peptide identification and quantification a library free search was performed using DIA-NN version 1.8.1 (Demichev *et al*. 2020) against the organism-specific Uniprot database (release 2023_03, *Synechocystis* sp. strain PCC 6803, taxid 1148), complemented with the sequences of RNase J, 3xFLAG and GFP. Trypsin was set as protease with 2 allowed missed cleavages. Carbamidomethylation of cysteine was selected as fixed modification, N-terminal excision of methionine and oxidation of methionine were selected as variable modifications with a maximum number of 2. For precursor ion generation, peptides ranging from 7 to 35 amino acids, precursor charge from 2 to 4, precursor *m/z* range from 361 to 1,100 and fragment ion *m/z* range from 200 to 1,200 were used. Further parameters were: match between runs, unrelated runs, search in double-pass mode, deactivated heuristic protein inference, robust LC quantification and RT-dependent cross-run normalisation. The protein group quantities output file from DIA-NN was used and all proteins with non-zero label-free quantification (LFQ) intensities in 3 out of 4 replicates in both groups were selected for further analysis. Missing values were imputed by sequential imputation using the R package impSeqRob (Verboven, Branden and Goos 2007). For differential abundance analysis, log_10_ transformed LFQ intensities were subjected to a moderated t-test using the R package limma (Ritchie *et al*. 2015) with correction of p-values for multiple testing using Benjamini and Hochberg’s method.

## Results

### Evidence that RNase J is involved in CRISPR3 crRNA maturation

We first tested the possible effects of RNase J on synthetic RNA substrates *in vitro.* We cloned and expressed histidine-tagged recombinant RNase J as described (Cavaiuolo *et al*. 2020). Purified RNase J was incubated with synthetic oligoribonucleotide RB27 corresponding to the CRISPR3 spacer 2-repeat sequence (substrate, 72 nt). Several weak bands were observed already in the absence of added enzyme indicating slight degradation of the substrate (background). However, a few bands clearly appeared or were increased in intensity upon addition of RNase J. Especially, the more prominent signal at 35 nt (**Fig. 1A**, lane 2), consistent with the primary cleavage at the end of the spacer as inferred from the analysis of primer extension products (**Fig. 2**), followed by exonucleolytic removal of single nucleotides close to the 5′ end of the repeat, as this would yield precisely a 35 nt fragment. Thus, RNase J may have processed the synthetic RNA via its 5′ exonuclease and endonuclease activities combined.

**Figure 1.**
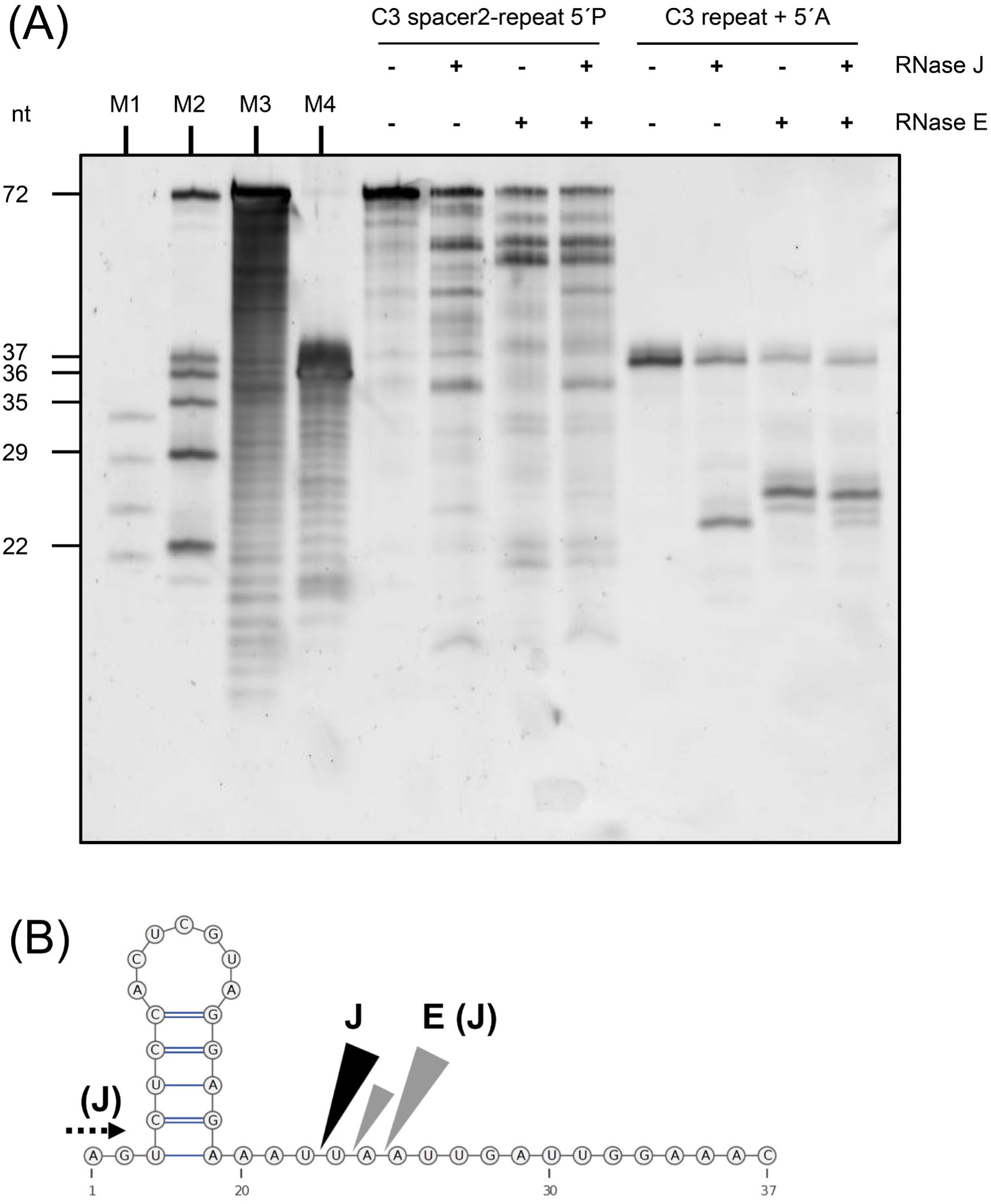
*In vitro* cleavage of CRISPR3 precursor RNA by RNase J. **(A)** Purified recombinant RNase J and RNase E (400 ng) were incubated in various combinations with 5 pmol of 5’-phosphorylated synthetic oligoribonucleotides as substrates. Lanes 1 to 4: Incubation with the 72 nt long CRISPR3 (C3) spacer 2-repeat RNA. Lanes 5 to 8: Incubation with the 37 nt long C3 repeat with an additional 5′A corresponding to the final nucleotide of spacer 2. Samples were incubated for 30 min at 30°C. The reaction was stopped by adding 2× RNA loading dye and denatured for 10 min at 70°C. Samples were analyzed on 8 M urea-15% PAA gel. The gel was stained with SyBR Gold (Invitrogen) in a 1:10,000 dilution with 0.5× Tris-Borate-EDTA buffer. Signals were detected with a Laser Scanner Typhoon FLA9500 (GE Healthcare) with the following settings: excitation, 473 nm; emission filter long pass blue ≥ 510 nm; photomultiplier value, 400. M1: ZR small RNA ladder (17, 21, 25, and 29 nt in length, Zymo research); M2: C3 ladder made of CRISPR3 repeat-related synthetic oligoribonucleotides of 72, 37, 36, 35, 29 and 22 nt in length (indicated on the left); M3: C3 spacer 2-repeat alkaline hydrolysis ladder; M4: C3 repeat + 5′A alkaline hydrolysis ladder; nt: nucleotide; C3: CRISPR3. Note that the marker bands do not run exactly according to size. **(B)** The CRISPR3 repeat RNA forms a hairpin at its 5′ end. Cleavage sites by RNase J (J, black wedge) and RNase E (E, gray wedges) are located within an adenylate–uridylate-rich sequence downstream of the stem-loop. The twin RNase E cleavage sites match previously determined sites 13 and 14 nt from the 3’ end (Behler *et al*. 2018). The 2 nt shorter fragment generated by RNase J compared to the RNase E main product suggests the cleavage site indicated by the black wedge. However, alternatively RNase J may have cleaved the same site as RNase E as an endoribonuclease, followed by 5’ exonucleolytic removal of 2 nt until reaching the repeat stem loop blocking further processing. This is indicated by the bracketed (J) and the dashed arrow. RNA structures were predicted by RNAfold (Hofacker 2003) and visualized using VARNA (Darty, Denise and Ponty 2009).

**Figure 2.**
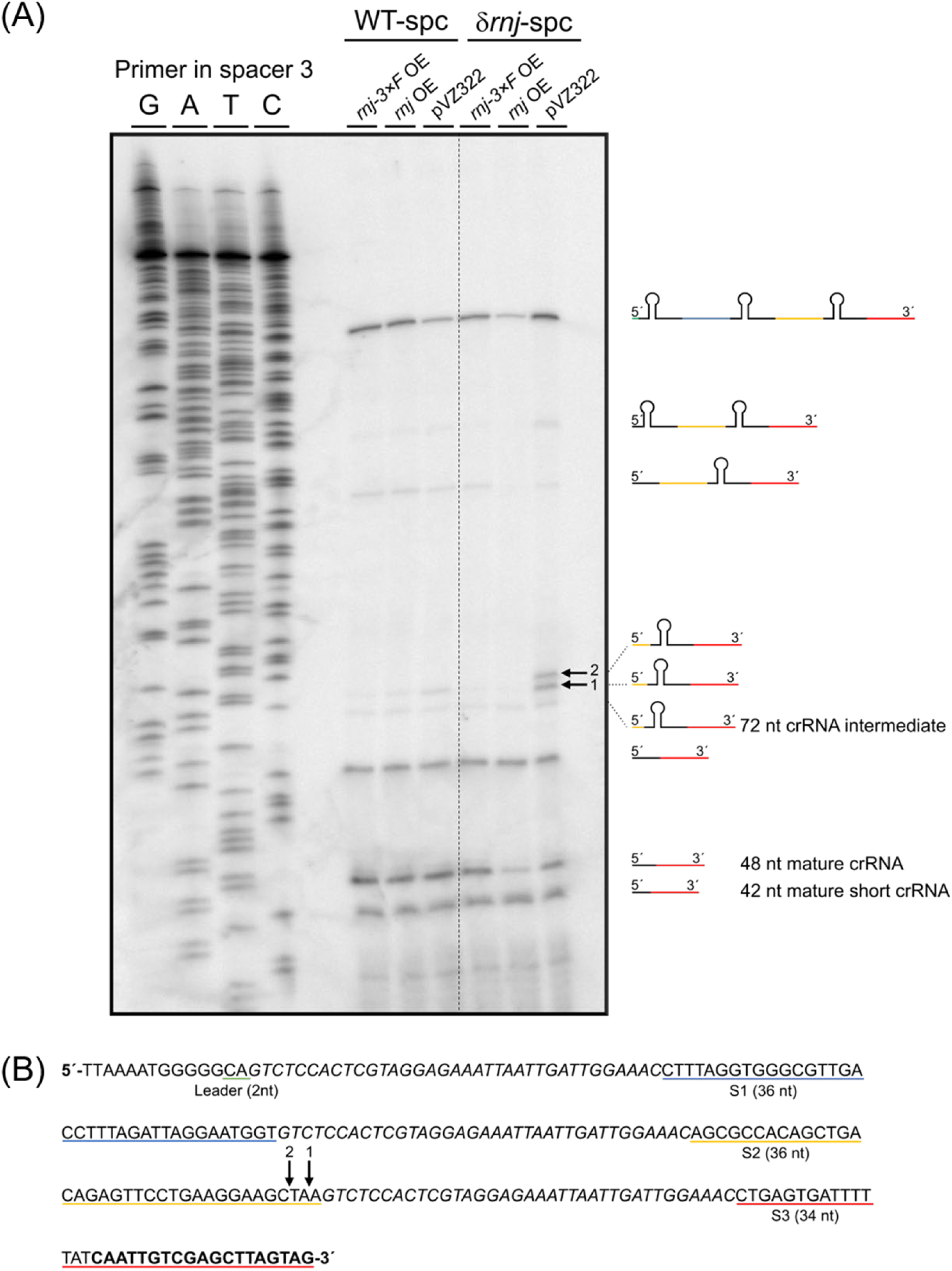
Mapping of RNase J *in vivo* processing sites within CRISPR3 pre-crRNA by primer extension. Total RNA was isolated from the *Synechocystis rnj* (*slr0551*) knock-down strain δ*rnj* carrying the empty vector pVZ322 as control, pVZ322-P*petE*-*slr0551*, or pVZ322-P*petE*-*slr0551-3×flag*, as well as the *Synechocystis* wild type with integrated spectinomycin cassette (WT-Spc) and carrying the same set of plasmid constructs as δ*rnj*. The RNA was subjected to primer extension analysis using the 5′-end-labelled oligonucleotide primer Ext_S3_rev (**Table S1**) and 2 µg of total RNA per reaction. Primer extension products were separated on 8.3 M urea-15% PAA gels cast in a sequencing gel device. **(A)** Mapping of *in vivo* cleavage within CRISPR3 revealed the stronger accumulation of two fragments in the δ*rnj* strain with decreased RNase J expression (pVZ322), labeled by arrows 1 and 2. **(B)** Section of the CRISPR3 repeat-spacer array nucleotide sequence, corresponding to the mapped region. The binding site of the labeled primer in spacer 3 necessary for reverse transcription is shown in bold. The sequence of the CRISPR3 repeats are in italics. The sequences of the spacers are underlined in blue for spacer 1 (S1), in yellow for spacer 2 (S2) and in red for spacer 3 (S3), with the length of the corresponding spacer in parentheses. The sequence of the 2 nt long leader is underlined in green. The cleavage sites attributed to RNase J are marked by black arrows labelled 1 for the +1 and 2 for the + 3 cleavage sites close to the end of spacer 2.

The same experiment performed with purified RNase E provided a different cleavage pattern (**Fig. 1A**, lane 3). A prominent appearing band likely is the substrate after 13 nt were cleaved off from the 3’ end. When incubating the RNA with both ribonucleases, we obtained a cleavage pattern corresponding to the two single digestions combined (lane 4). Because the appearance of multiple bands complicates the interpretation, we then performed analogous experiments with a single synthetic CRISPR3 repeat sequence with a single additional adenosine at the 5′ end yielding a 37 nt substrate. Incubation with RNase E alone resulted in one major band and a minor band one nt shorter (**Fig. 1A**, lane 7). These fragments match previous observations made in *in vitro* assays using synthetic CRISPR3 repeat sequences as substrate (Behler *et al*. 2018) and release the unusually long, 13 and 14 nt long 5’ handles observed *in vivo* (Scholz *et al*. 2013). Incubation with RNase J alone resulted in single major band as well (**Fig. 1A**, lane 6), with a size difference of 2 nt compared to the digestion with RNase E (**Fig. 1A**, lanes 6 and 7). The RNase J cleavage site thus must be close to the processing site used by RNase E to generate the 5′ handle of the CRISPR3 crRNA (**Fig. 1B**). Alternatively, RNase J cleaved the same site as RNase E as endoribonuclease, followed by by exonucleolytic removal of 2 nt until reaching the repeat stem loop blocking further processing (**Fig. 1B**).

The absence of a ladder-like pattern suggested that primarily the endonuclease activity of RNase J was active. When mixing both enzymes, we obtained a cleavage pattern qualitatively corresponding to the two single digestions combined, but quantitatively, the RNase J-assigned signal was weaker (**Fig. 1A**, lane 8). This result matches the observation that *in vivo* RNase E is the dominating and rate-limiting enzyme in processing the CRISPR repeat (Behler *et al*. 2018). The *in vitro* cleavage data in **Fig. 1** provide critical evidence that RNase J processes CRISPR3 RNA precursors.

### RNase J expression influences processing of crRNA intermediates *in vivo*

To determine whether RNase J affects the maturation of CRISPR3 crRNA *in vivo*, primer extension analysis was performed to map possible processing differences between the strains overexpressing tagged and untagged RNase J *in vivo*. Therefore, we cloned the *slr0551* gene encoding RNase J into the pVZ322 vector under the control of the copper-inducible *petE* promoter, P*_petE_* (Zhang *et al*. 1992). We designed two different constructs: one expressing the unmodified RNase J protein, and one expressing a version with a 3×FLAG sequence fused to its C-terminal end. The constructed plasmids were conjugated into *Synechocystis* strain PCC-M (Trautmann *et al*. 2012) via tri-parental mating (Scholz *et al*. 2013). In addition, we included the *Synechocystis* δ*rnj* strain with a reduction in the number of *rnj* loci per cell (partial gene deletion) (Cavaiuolo *et al*. 2020), in which we overexpressed C-terminally tagged, or untagged RNase J. The *rnj* overexpression in *Synechocystis* WT-spc and δ*rnj*-spc strains carrying pVZ322-P*_petE_*-*rnj* compared to the control strains with the empty pVZ322 vector as control was verified by northern hybridization (**Supplemental Fig. S1**). Because the partial deletion of *rnj* gene copies by insertion of a streptomycin resistance gene in the *Synechocystis* δ*rnj* strain depends on the presence of streptomycin, we used as background for all engineered strains in this experiment not the wild type, but strain WT-Spc, with a spectinomycin cassette integrated into a neutral site within the *ssl0452-slr0271* intergenic region (Cavaiuolo *et al*. 2020). A strain carrying the empty pVZ322 vector was used as control. The cells were cultivated until reaching an OD_750_ of 0.8 and cells were collected 24 h after RNase J expression was induced. Primer extension was performed on 2 µg of total RNA using the same primer previously used to analyze the involvement of RNase E in CRISPR3 maturation (Behler *et al*. 2018). The result is shown in **Fig. 2**. The WT-Spc control strain yielded an identical pattern to previous primer extensions obtained with the *Synechocystis* wild type (Behler *et al*. 2018), indicating that the integrated resistance cassette had no effect on crRNA processing. Moreover, the two strains overexpressing RNase J or RNase J-3×FLAG showed identical patterns to the WT-Spc strain. However, two differences were apparent in the δ*rnj* strain, harboring only the empty vector. In this strain, one additional band and one band with increased intensity were observed (**Fig. 2**, arrows). While the lower band (arrow 1) was weaker, but present also in the other lanes, the upper band (arrow 2) was unique to the δ*rnj* strain (**Fig. 2**). Because in this strain the RNase J level is the lowest, this observation indicated that

a. the reduced level of RNase J in the δ*rnj* strain impacted the processing of the CRISPR3 crRNA maturation and
b. this effect was reversed when overexpressing recombinant RNase J from a plasmid.

Using the sequence ladder on the left, we identified the corresponding sequences of the observed bands. The 3’ ends of the two fragments were located close, 1 and 3 nt, to the 3’ ends of spacer 2 (S2) of the CRISPR3 array (**Fig. 2B**, arrows 1 and 2). As these two fragments accumulated in the cell, we conclude that the depletion in RNase J caused a bottleneck in the processing of the crRNA.

Previous work identified the first cleavage site of the CRISPR transcript is located at the end of the spacer, at exactly these positions, 1 to 3 nt from the first nt in the repeat (**Fig. 2B**, arrows), leading to the 72 nt intermediate (see Fig. 3 in Scholz *et al*. (2013)). Thus, the corresponding transcript was a substrate for RNase J activity, which was limiting in this strain. As RNase J from *Synechocystis* possesses 5′ exonuclease activity (Cavaiuolo *et al*. 2020), it may then remove 2 to 4 further nucleotides from the 5′ end until reaching the stem-loop structure beginning at nt 2 of the repeat. That structure then is recognized by RNase E in a further processing step, which generates the crRNA 5′ handle (Behler *et al*. 2018).

**Figure 3.**
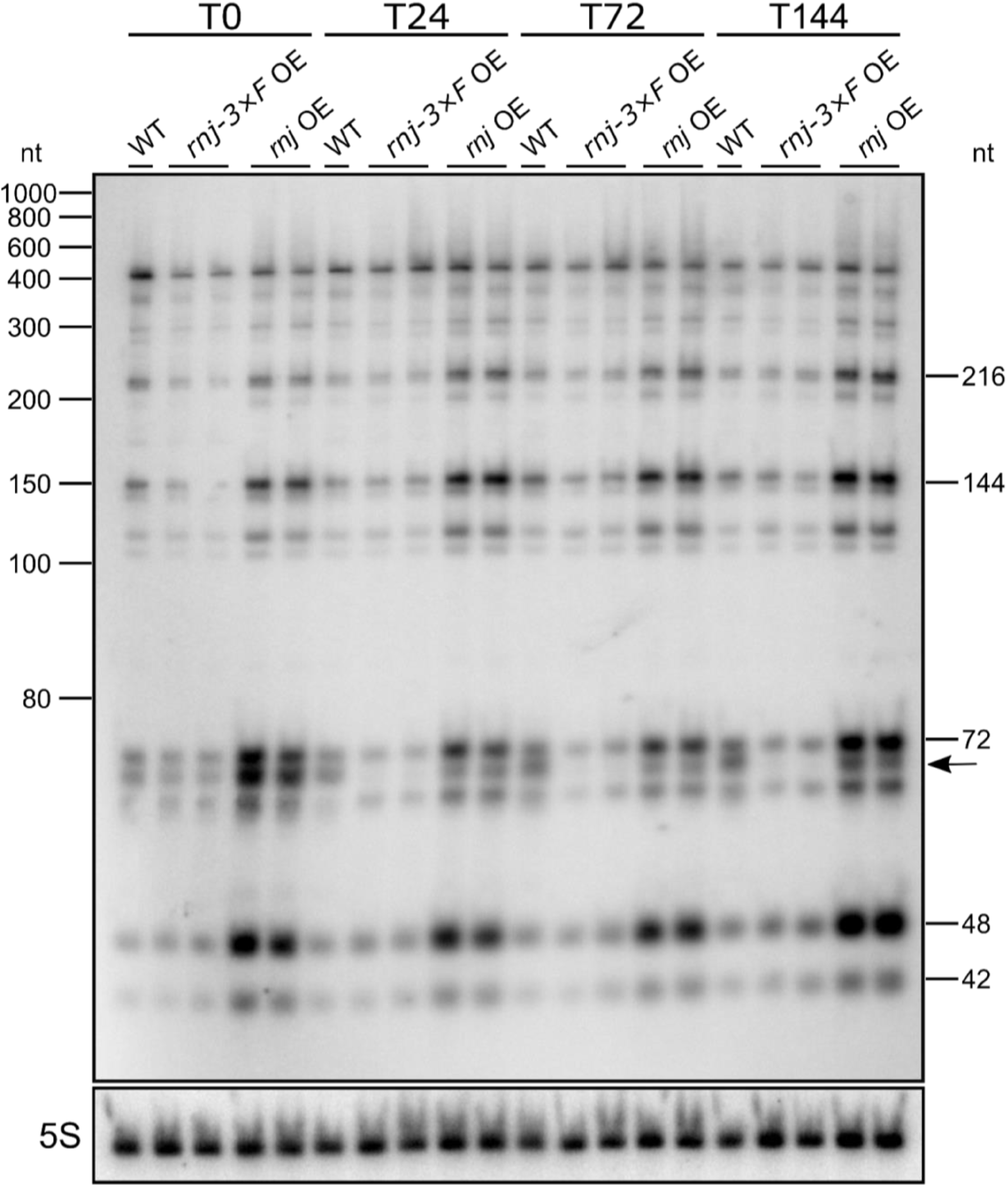
CRISPR 3 crRNA accumulation in *Synechocystis* wild type and RNase J overexpression strains. The strains carrying pVZ322-P*_petE_* constructs for the expression of RNase J (*rnj* OE) or RNase J-3×FLAG (*rnj* x3F OE) and the wild type strain (WT) were cultivated in copper-free BG11 medium until reaching an OD_750_ of 0.8. Expression of RNase J was induced with 2 µM CuSO_4_. Samples of the cultures were collected before induction (T0), and 24 h, 72 h and 144 h after induction. Ten μg of total RNA isolated from each culture was separated on an 8.3M urea - 10% PAA gel. For Northern hybridization, a ^32^P-labelled transcript probe was used spanning CRISPR 3 spacers 1 - 4. Hybridization of 5S rRNA is shown as the control for equal loading. The black arrow marks the band at approximately 70 nt that is referred to in the text. The *rnj* overexpression was verified by northern hybridization (**Supplemental Fig. S1**).

Northern analyses were used to validate possible effects of RNase J overexpression on the accumulation of CRISPR3 processing intermediates. *Synechocystis* wild type served as control. Samples were harvested over a time course of 24, 72 and 144 h post induction and RNA was isolated for each strain. The impact of ribonuclease J overexpression on CRISPR3 crRNA maturation was analyzed using a radioactively labeled single-stranded RNA probe spanning the CRISPR3 spacers 1 to 4. The resulting complex pattern of mature crRNAs and processing intermediates in the wild type was fully consistent with previous analyses (Scholz *et al*. 2013; Behler *et al*. 2018). The mature crRNA forms of 48 and 42 nt were among the strongest signals (**Fig. 3**). Another strong signal was obtained for the 72 nt, 144 nt and 216 nt long processing intermediates consisting of single, double and triple repeat-spacer units. Moreover, the previously reported 6 nt-periodicity (Behler *et al*. 2018) was visible in the form of the double bands for the mature crRNAs and all processing intermediates. At time point T0, prior to induction, we expected that the wild type and the two overexpression strains should show quantitatively and qualitatively similar signals. While all strains did show similar patterns, most band intensities were stronger in the strain carrying the construct for the overexpression of the untagged enzyme (*rnj* OE). Especially striking was the difference for the smaller RNA fragments, the 144 nt and shorter processing intermediates, and the mature crRNAs. This observation may be explained by the leakiness of the P*_petE_* promoter. Furthermore, we noticed a slight reduction in the intensities of the signals between 100 and 250 nt in the strain encoding the tagged enzyme (*rnj-3xF* OE) (**Fig. 3**). This difference in activity between the tagged and untagged RNase J became more pronounced during the later times of the time course.

The most striking difference was seen in the disappearance of a ∼70 nt band (marked by a black arrow in **Fig. 3**) in the strain overexpressing the tagged enzyme after induction. The size of the band slightly below the main signal for the single repeat-spacer unit of 72 nt indicates that it is an crRNA maturation intermediate 2 or 3 nt shorter. The increased signal intensities in the strain overexpressing untagged RNase J confirmed that RNase J is involved in the maturation of crRNA from the subtype III-Bv CRISPR system in *Synechocystis*. Furthermore, the tagging of RNase J at its C-terminal end may have enhanced its exoribonuclease activity, explaining the overall lower crRNA signal intensities in this strain and especially the disappearance of the signal at ∼70 nt at the T24, T72 and T144 time points in the strain overexpressing the tagged enzyme. Previous observations indicated an enhanced 5’ exoribonuclease activity for C-terminally poly-histidine-tagged RNase J1 in *Bacillus subtilis* (Laalami and Putzer 2011), which would be consistent with the observations made here for the C-terminally tagged RNase J.

### Characterization of proteins interacting with RNase J

We next sought to identify potential interaction partners of RNase J. Co-IP experiments were performed with *Synechocystis* strains expressing RNase J either N- or C-terminally tagged with the 3×FLAG epitope, and with GFP-expressing strains as controls (plasmids pVZ322-P_rha_-*3×FLAG*-*slr0551*, pVZ322-P_rha_-*gfp*-*3×FLAG*, pVZ322-P*_petE_*-*slr0551*-*3×FLAG,* or pVZ322-P*_petE_*-*gfp*-*3×FLAG*). The strains were cultivated in quadruplicates in high density cultivation systems (CellDEG GmbH) until an OD_750_ of approximately 9–12 was reached, and protein expression was induced for 24 h before harvesting the cells, with a final OD_750_ of approximately 25–30. Triple-FLAG tagged proteins were purified with M2 anti-FLAG magnetic beads and analyzed by SDS-PAGE (**Supplemental Fig. S2 and S3**). Potential interaction partners were identified by mass spectrometry. In the entire dataset, a total of 2,625 proteins was identified, corresponding to ∼71.5% of the total proteome. The list of proteins identified for each replicate is detailed in **Supplemental Table S2**.

In order to identify potential interaction partners, a log_2_FC threshold of +1.5 (p value ≤ 0.05) was established to select proteins which were at least 2.8-fold enriched in the RNase J overexpression strains with respect to the GFP expressing control strains. With this parameter, 20 proteins were identified for the N-terminal tagged (NTT) RNase J and 36 proteins for the C-terminal FLAG-tagged (CTT) RNase J. In both cases, RNase J, used as bait, showed the highest respective enrichment (log_2_FC of +10.26 and +12.13). In addition to RNase J, eight enriched proteins were shared between the two co-IPs with the differentially tagged RNase J variants.

A gene ontology analysis was conducted to categorize the identified proteins according to their PANTHER protein class (Thomas *et al*. 2022). Approximately one third of the detected proteins were not assigned to a PANTHER category (35.0% for NTT and 35.3% for CTT RNase J). In both batches, the second most enriched prominent group (after unclassified) was that of metabolite interconversion enzymes (35.00% in NTT and 14.70% in CTT RNase J). In CTT, proteins were assigned to DNA metabolism (11.80%), protein modifying enzymes (8.80%) and translational proteins (5.90%) which were not found in the NTT batch. Chaperone proteins represent 10.0% in the NTT batch and 5.9% in the CTT batch (**Supplementary Table S3**).

The calculated log_2_FC data were plotted over the adjusted p values in the form of a volcano plot (**Fig. 4**). Among the nine proteins shared between the co-IPs with the two differentially tagged RNase J variants, PNPase stood out with log_2_FC enrichment factors of +4.67 and +5.96 (**Table 1**). Five of the other proteins that were found consistently enriched with both NTT and CTT RNase J have regulatory functions. These were EbsA, a factor essential for biofilm self-suppression that interacts with Hfq and PilB (Yegorov *et al*. 2021), the PatA family regulator of phototaxis PisG (Yoshihara *et al*. 2000), the RuBisCO transcriptional regulator (Bolay *et al*. 2022), Sll0245, a possible homolog the ribosome-binding ATPase YchF (Tomar, Kumar and Prakash 2011), and Sll0086, annotated as a one-domain ArsA-like protein, although experimental analyses indicated that it is not involved in arsenic or antimony resistance (López-Maury, Florencio and Reyes 2003). Further consistently enriched proteins were Slr0244 containing a universal stress protein A (UspA) domain, and Slr0552, an unknown protein encoded by a gene directly downstream of *rnj* (**Table 1**).

**Figure 4.**
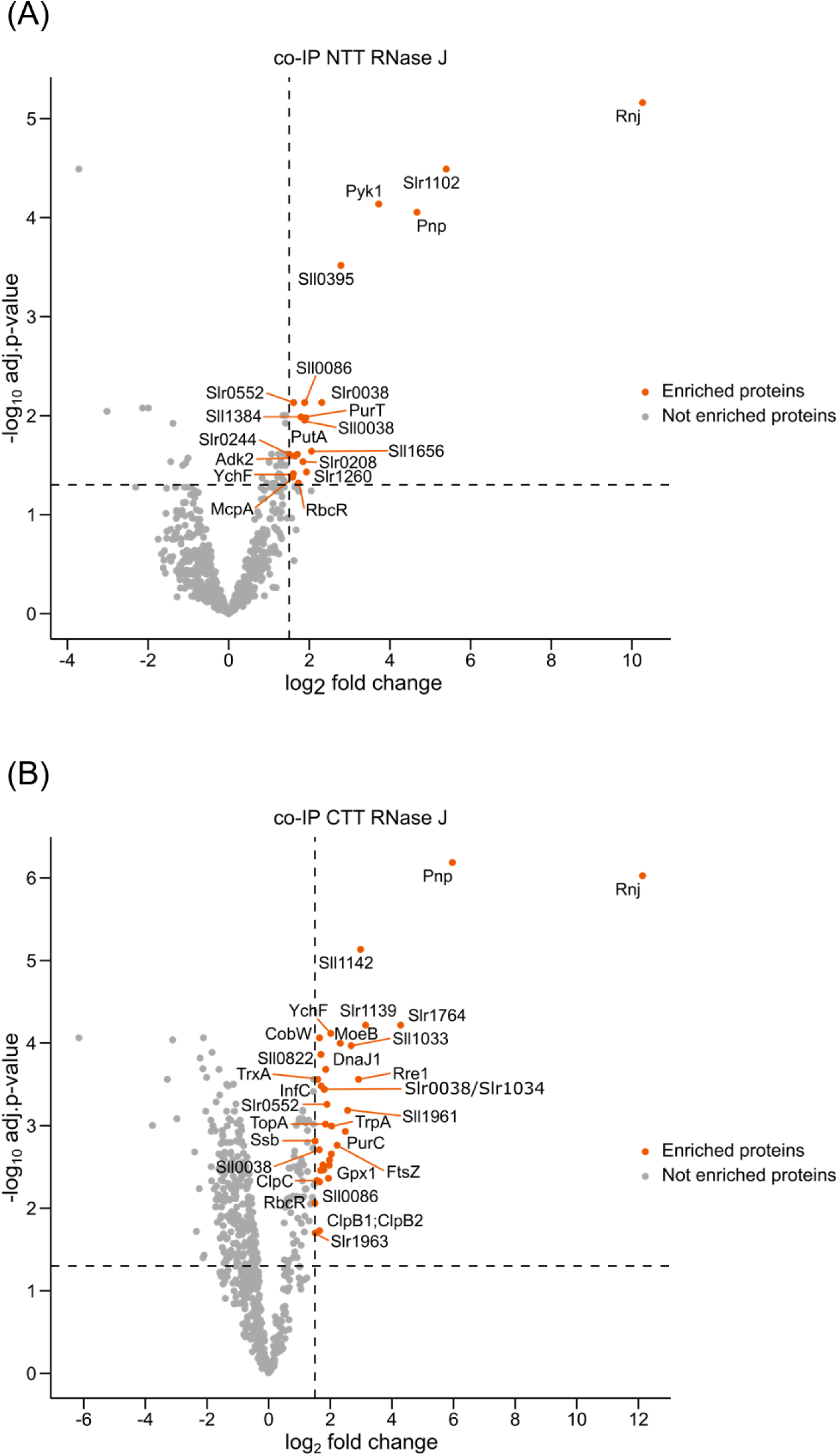
Co-IP experiment for the identification of interaction partners of RNase J. *Synechocystis* strains carrying pVZ322 constructs for the expression of RNase J with N-or C-terminal 3×FLAG tag, under the control of rhamnose or *petE* promoter, respectively, were cultivated in the CellDEG high density system. *Synechocystis* strains overexpressing GFP-3×FLAG under the control of the same promoters were used as control. Protein expression was induced after 48 h of cultivation with 6 mg/ml L-rhamnose or 2 µM CuSO_4_. The equivalent of 1000 OD cells was harvested. FLAG-tagged proteins were purified with M2 anti-FLAG magnetic beads and co-enriched proteins were analyzed by mass spectrometry. Proteins detected in the RNase J elution fraction were compared with the corresponding GFP control and visualized as volcano plots based on a Limma moderated t-test with adjustment for multiple testing correction. The thresholds for p value and log_2_ fold change (log_2_FC) were set to 0.05 and 1.5, respectively. Proteins with significant enrichment are indicated by orange dots (enriched), while proteins which did not match the selection criteria are shown in light grey (not enriched). **(A)** Co-IP with NTT RNase J. See **Supplemental Fig. S2** for the Coomassie-stained SDS-PAA gel. **(B)** Co-IP with CTT RNase J. See **Supplemental Fig. S3** for the Coomassie-stained SDS-PAA gel. Details of the proteins consistently enriched in both co-IP experiments are provided in **Table 1**. The complete mass spectrometry data set is provided in **Table S2**.

**Table 1.**
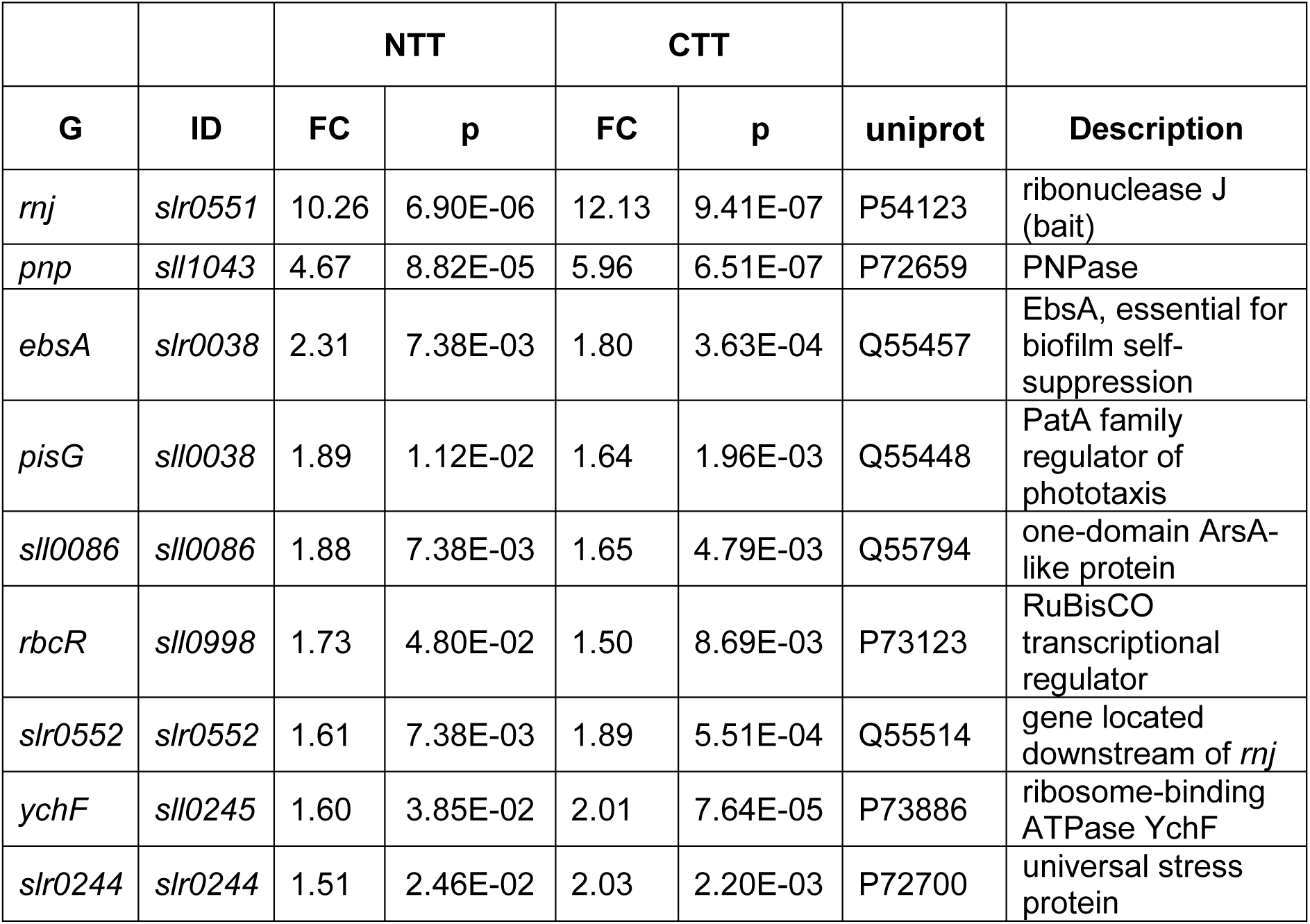
List of proteins enriched in the co-IP with RNase J. These are the proteins enriched with both, N-terminally 3×FLAG-tagged (NTT) and C-terminally 3×FLAG-tagged (CTT) RNase J and passing the selection threshold (log_2_FC ≥1.5, p value ≤0.05). In addition, these proteins were detected in at least three out of four replicates with NTT and CTT RNase J. The columns indicate gene (G) and locus IDs, followed by the log_2_FC (FC) and p values (p) for the separate enrichments, uniprot accessions and descriptions. Results from the MS analysis are also plotted in **Fig. 4**.

### PNPase is not involved in CRISPR3 crRNA maturation

PNPase was the most highly enriched ribonuclease in the RNase J co-IP (**Fig. 4A, B**). Moreover, the PNPase in *Synechocystis* forms a complex with RNase E (Zhang *et al*. 2014), therefore it was considered as another candidate for participation in the CRISPR3 crRNA maturation in *Synechocystis*. We expressed PNPase as a recombinant protein in *E. coli*, purified and tested its activity in *in vitro* cleavage assays. The samples with the successive addition of PNPase or RNase E showed the 22 and 23 nt cleavage products expected for cleavage by RNase E and remaining substrate (Behler *et al*. 2018). In all samples in which PNPase was added first, the intensity of the 36 nt substrate was lowered indicating the rapid degradation of the substrate (**Fig. 5**). Moreover, these samples showed a single nt ladder although of very low intensity, directly below the substrate. When RNase E was added first, all samples showed a stronger intensity of the 22 and the 23 nt products, likely because the entire substrate amount had been available for the cleavage by RNase E or because RNase E, when bound to the RNA, protects the substrate from the 3′exonuclease activity of PNPase. The other way around, when PNPase was added first, a part of the substrate was degraded first and only the rest of the substrate was available for the cleavage by RNase E (**Fig. 5**).

**Figure 5.**
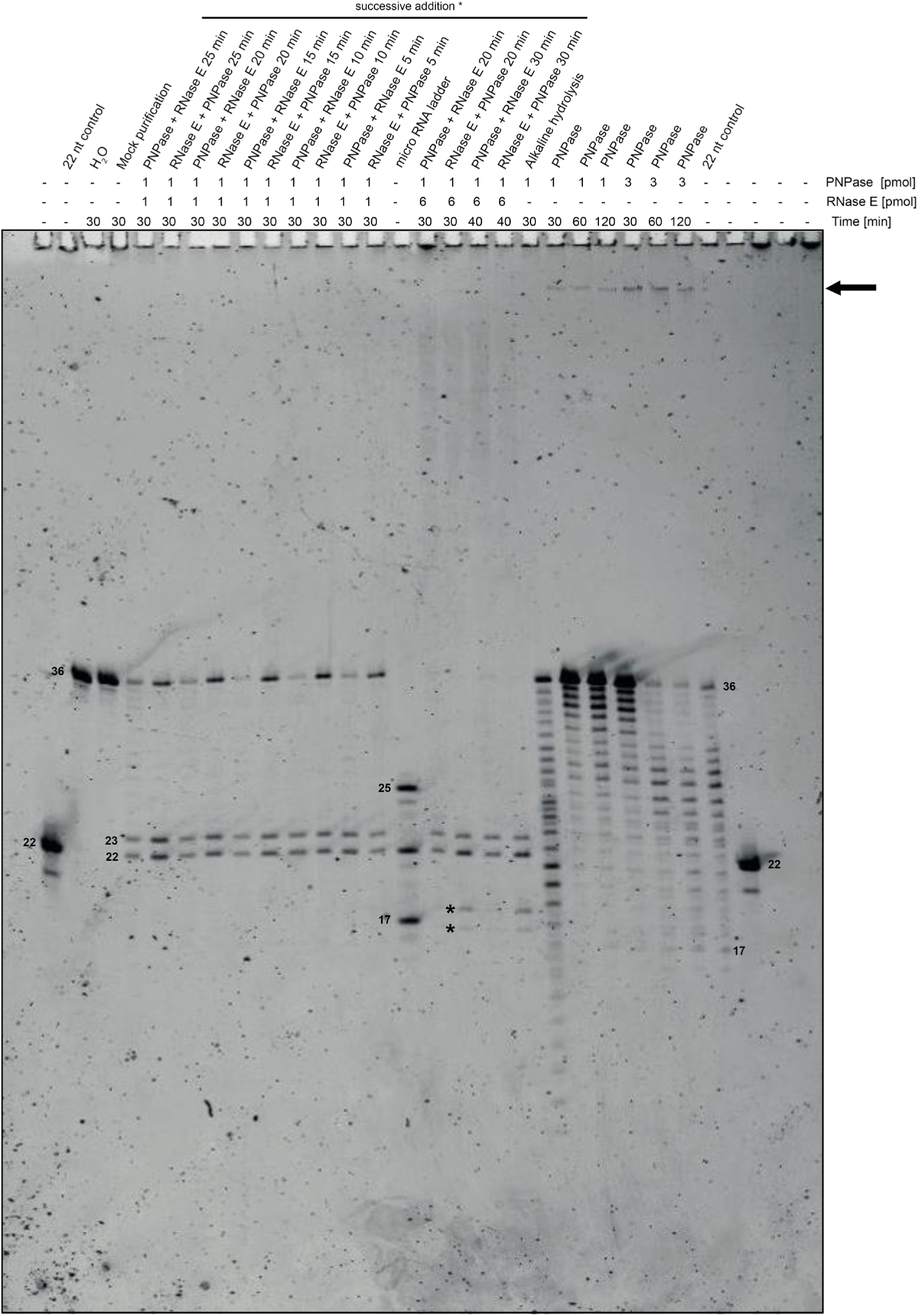
*In vitro* cleavage assays with PNPase and RNase. **E.** Different concentrations of each enzyme were used. The 5’-P CRISPR3 repeat RNA oligonucleotide was used as substrate (100 ng per reaction) at 30°C with the indicated enzyme amounts and incubation times. The first enzyme was added directly at the beginning and the second enzyme was added after the time indicated behind. 8 M urea - 15% PAA gels were used and stained with SYBR Gold. The Micro RNA ladder (NEB) served as size marker. All sizes given in nt. Nuclease-free water and the mock purification served as negative controls. A black arrow refers to unusual high molecular products. Asterisks label unspecific products resulting from high RNase E concentrations if this enzyme was added first.

Samples incubated only with PNPase, irrespectively of incubation times and whether 1 or 3 pmol enzyme were used, showed a single nt ladder, although more substrate was degraded when more PNPase was used (**Fig. 5**, lanes to the right). We noticed that the ladder pattern was restricted to the 17 to 36 nt size range. This limitation was not an effect of the used gel conditions as the alkaline hydrolysis of the same substrate showed the ladder extending much further down. However, this result fits to the secondary structure of the CRISPR repeat used as substrate: it consists of a single stem loop from position 2 to 18, while the sequence from nt 19 to 36 is single-stranded (Behler *et al*. 2018). Hence, this result is consistent with a strong 3’ exonuclease activity of the *Synechocystis* PNPase that stopped once the stem loop was reached.

## Discussion

### RNase J is involved in the CRISPR3 crRNA maturation

In the type III-Bv CRISPR3 system of *Synechocystis*, RNase E replaces the lacking Cas6 activity in the maturation of the pre-crRNA (Behler *et al*. 2018). Although RNase E is responsible for a major processing step, it has been suggested that additional ribonucleases are involved. While RNase E cleaves within the CRISPR3 repeat RNA, directly generating the characteristic 13 nt and 14 nt 5′ repeat handles (Behler *et al*. 2018), it does not generate the 3’ ends of the mature crRNAs. Based on the analysis of mapped transcriptome reads, we previously characterized CRISPR3 crRNA maturation as a 2-step process (Scholz *et al*. 2013). This analysis revealed that the first cleavage site is located in the spacer and not in the repeat. This conclusion was supported by the observation that the 72 nt processing intermediate spans across the cleavage site within the repeats and ends at the cleavage site in the spacer (Scholz *et al*. 2013).

Here, we show that overexpression of RNase J correlates with an increased abundance of the ∼72 nt fragment, which represents a single repeat-spacer unit, of other intermediates, and of the 48 nt and 42 nt mature crRNAs (**Fig. 1**). The results of the primer extensions clearly identified two 5’ ends that were enriched in the δ*rnj* strain with decreased RNase J activity (**Fig. 2**). These results, in conjunction with the results of Behler et al. (2018), suggest that RNase J mediates 5’ exonucleolysis immediately after the first cleavage, an endonucleolytic cleavage close to the spacer 3’ ends. After this, RNase E releases the mature crRNA 5’ ends including the 13 or 14 nt 5′ repeat handle (the 1 nt size difference in the length of the repeat handle likely results from ambiguity in the substrate cleavage by RNase E itself (Behler *et al*. 2018)).

Looking for proteins that interact with RNase J could reveal factors that potentially influence the activity or substrate recognition of RNase J, or act as an additional nuclease involved in further 3’ processing. Such processing is required as two mature crRNA forms accumulate *in vivo*, 42 and 48 nt in length and only the latter results from the combined action of RNase E and J. Therefore, we searched for proteins possibly interacting with RNase J. This search identified PNPase as an enzyme strongly co-enriched with RNase J (**Fig. 4**). PNPase is participating in crRNA maturation in *Listeria monocytogenes* (Sesto *et al*. 2014) and *Bacillus subtilis* (Chou-Zheng and Hatoum-Aslan 2019). In the Type III-A CRISPR–Cas system of *Staphylococcus epidermidis*, Csm5 was found to interact and stimulate PNPase activity (Walker *et al*. 2017). These findings prompted us to overexpress and test PNPase from *Synechocystis*. While the characterization of *Synechocystis* PNPase in an *in vitro* assay demonstrated a strong 3’ exonuclease activity (**Fig. 5**), we found no evidence supporting its involvement in CRISPR3 maturation, at least *in vitro*. In the elution of the CTT RNase J, the bifunctional nuclease domain protein Sll1142 was identified. Homologs of this protein are conserved in cyanobacteria, and related proteins exist in many other bacteria and plants. A homolog in *Oryza minuta* was reported to possess bifunctional RNase and DNase activities *in vitro* (You *et al*. 2010). However, whether Sll1142 has any CRISPR-related function remains to be elucidated.

The here described involvement of RNase J in the maturation of CRISPR3 crRNAs adds further evidence on the connection between degradosome components and the type III CRISPR-Cas machinery (Behler *et al*. 2018; Chou-Zheng and Hatoum-Aslan 2019; Bilger *et al*. 2024). Our current model for CRISPR3 crRNA maturation is shown in **Fig. 6**.

**Figure 6.**
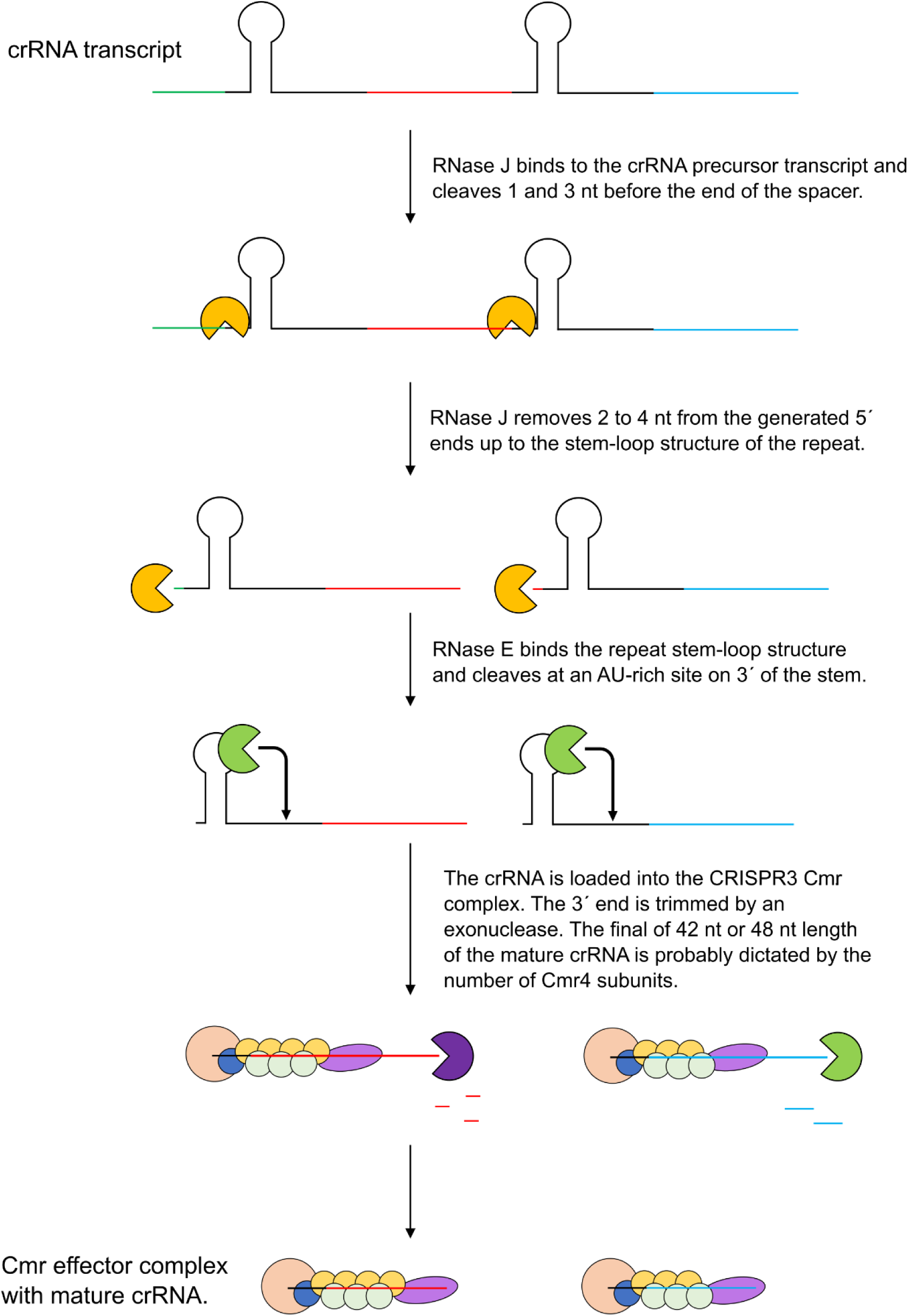
Model for the initiation of the maturation of CRISPR3-derived crRNAs. The transcribed CRISPR3 pre-crRNA (repeats in black) is recognized by RNase J (yellow) cleaving the transcript 1 and 3 nt from the spacer 3’ end, described as the initial processing step (Scholz *et al*. 2013), followed by 5′end trimming by 2 to 4 nt until reaching the stem-loop structure located at the beginning of the repeat sequence. RNase E (green) then recognizes the CRISPR repeat stem-loop structure and cleaves the RNA in the AU rich region downstream of the stem. This yields the mature 48 nt long crRNAs with a 13 or 14 nt 5′handle (Behler *et al*. 2018). The pre-crRNA is then loaded into the Cmr complex. Further 3′ trimming to the 42 nt mature crRNAs requires an unknown 3′ exoribonuclease (purple). The final length of 42 nt or 48 nt (Scholz *et al*. 2013) may be dictated by the number of Cmr4 subunits (yellow disks) attached to the crRNA (Hatoum-Aslan *et al*. 2013). Both the 42 nt and 48 nt mature crRNA forms accumulate *in vivo* (Scholz *et al*. 2013). The mature crRNAs are then loaded into the CRISPR3-Cas protein complex.

### Co-IP analysis indicates the existence of a degradosome in *Synechocystis* **involving RNase J**

We identified several potential interaction partners for the bifunctional endo- and exoribonuclease RNase J in *Synechocystis*. We conducted a co-IP experiment with N- or C-terminal 3×FLAG-tagged RNase J. The results revealed a significant difference between the two pull-downs. In addition to RNase J, 35 enriched proteins were identified for the CTT RNase J and 19 proteins for the NTT RNase J. Notably, only eight proteins were enriched in both co-IPs (**Table 1**). This discrepancy suggests that the presence and position of the 3×FLAG tag on RNase J was a relevant factor. The detection of five proteins with regulatory functions, EbsA, PisG, Sll0086, RbcR and YchF, that were consistently enriched in the co-IP with RNase J suggests an involvement of RNase J in regulatory responses that require ribonucleases to degrade transcripts that are no longer required.

Intriguingly, the most enriched protein together with RNase J was PNPase. The composition of RNA degradosomes varies considerably between different species (Tejada-Arranz, de Crecy-Lagard and de Reuse 2020; Zhang, Hess and Zhang 2022), but they share the presence of at least one major RNase (either RNase E, J or Y) and an RNA helicase (Tejada-Arranz, de Crecy-Lagard and de Reuse 2020). In the cyanobacteria *Nostoc* (*Anabaena*) sp. PCC 7120 and *Synechocystis*, RNase E forms a complex with PNPase, wherein the latter recognizes a conserved nonapeptide RRRRRRSSA at the end of RNase E (RRRRRSSA in *Synechocystis*) (Zhang *et al*. 2014). In our study, RNase E was not enriched in comparison to the GFP control. However, PNPase was the most enriched protein in both, CTT and NTT RNase J co-IPs. These data suggest that PNPase can interact not only with RNase E, but also with RNase J. RNase J carries a conserved arginine-rich motif close to its C terminus (RRSRKRS in *Synechocystis*), as is found in RNase E. Structure prediction using Alphafold 3 (Abramson *et al*. 2024) rendered five structures, four of which showed a potential interaction between the arginine-rich motif of RNase J and PNPase, although with only moderate pTM and iPTM support scores of 0.52 and 0.51, respectively (**Supplemental Fig. S4**). The residues that possibly bind PNPase to the C-terminal domain (CTD) of RNase J were compared with previously identified interaction sites of PNPase and RNase E in *E. coli* (Durán-Figueroa *et al*. 2006). The residues from our prediction were all located within the interaction domain of PNPase (**Supplemental Fig. S5**), indicating that the same region might be responsible for the interaction, though *E. coli* RNase E does not possess the arginine-rich motif from *Synechocystis* RNase E or RNase J. In *Synechocystis,* PNPase may form a complex with either RNase E or RNase J, but it is not possible to conclude which interaction is favored under standard growth conditions, as the overexpression of RNase J in the cell may have favored the binding of PNPase to ribonuclease J. In *M. tuberculosis*, PNPase, RNase E, RNase J, and RNA helicase RhlE were predicted to interact (Płociński *et al*. 2019). Similarly, in *Bacillus subtilis* RNase Y, PNPase, the helicase CshA and the glycolytic enzymes enolase and PfkA interact (Lehnik-Habrink *et al*. 2010). These studies have shown that the composition of the enzyme complexes that degrade RNA and the interactions among the involved enzymes are also rapidly changing (Cascante-Estepa, Gunka and Stülke 2016). In analogy, it cannot be excluded that *Synechocystis* RNase E and RNase J may interact together, or separately with PNPase, or other factors to degrade RNA.

While we identified RNase J as part of the machinery enabling the maturation of CRISPR3 crRNA in the absence of Cas6 enzymes, we were unable to identify enriched Cas proteins from the CRISPR3 effector complex. This indicates that RNase J does not perform an interaction with the effector complex, or that the crRNA is loaded into the Cmr effector complex at a later stage.

One uncharacterized protein that was enriched in both RNase J co-IPs was Slr0552. This is of interest because RNase J (Slr0551) and Slr0552 are encoded by two genes in the same operon. A BlastP search revealed that homologs exist in cyanobacteria; however, none of the homologs has been characterized so far.. Synteny analysis of the genomic organization of *rnj* and *slr0552* homologs show a conserved synteny in the *Synechocystis* genus and also in more distantly related cyanobacteria like *Gloeothece* and *Crocosphaera* (**Supplementary Fig. S6)**. Structural analysis with Foldseek and HHpred did not return any reliable results. Therefore, the function of Slr0552 and its relationship to RNase J remains a subject for future investigation.

## Data availability

MS proteomics data have been deposited to the ProteomeXchange Consortium via the PRIDE partner repository (Perez-Riverol *et al*. 2025) and are available through the identifier PXD060866.

## Acknowledgments

We thank Viktoria Reimann, Freiburg, for the 5S rRNA hybridization in **Fig. 3** and Juliane Behler, Freiburg, for helpful comments in an early phase of the project.

## Author contributions

WRH designed the project and secured funding. FD, BK and PH carried out MS-based proteomic analyses. TB carried out all PNPase-related experiments, HL contributed to the co-IP analyses, all other experiments were performed by RB. Data were analyzed by RB, FD, HP and WRH; RB and WRH wrote the manuscript with input from all authors.

