## Supplemental Information for "Involvement of RNase J in CRISPR RNA maturation in the cyanobacterium *Synechocystis* sp. PCC 6803"

### Supplementary Information

**Supplementary Figures:** p. 2

**Supplementary Tables:** p. 8

**Supplementary Methods:** p. 11

**Supplementary References:** p. 12

### Supplementary Figures

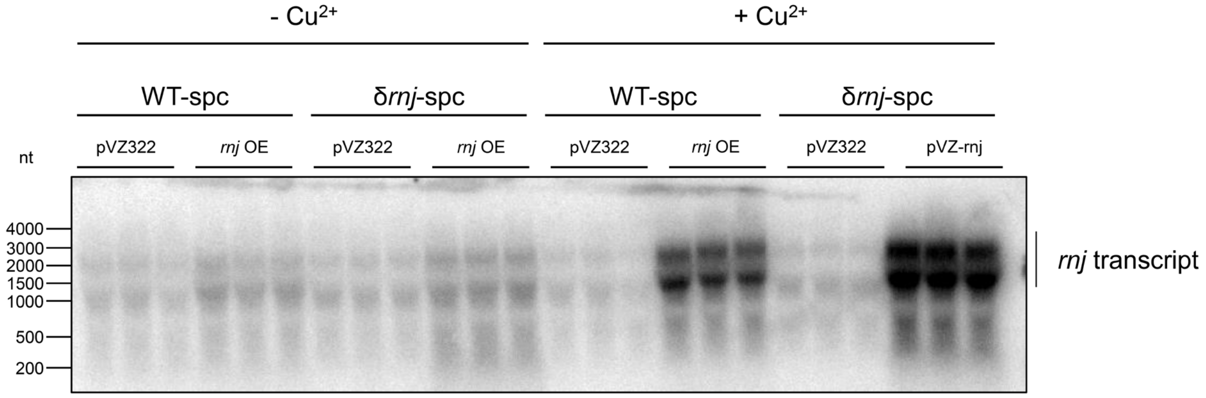


**Figure S1. Accumulation of *rnj* transcript in *Synechocystis* WT-spc and δrnj-spc strains carrying either pVZ322-P$_{petE}$-*rnj* or the empty pVZ322 vector as control.** The strains were cultivated in copper-free BG11 medium until reaching an $OD_{750}$ of 0.6. Expression of RNase J was induced with 2 µM $CuSO_4$. Uninduced cells served as control. 15 µg of total RNA were separated by electrophoresis in a 0.75% agarose-formaldehyde gel. For Northern hybridization, a $^{32}P$-labelled transcript probe was used targeting the *rnj* mRNA. The RiboRuler High Range RNA Ladder (Thermo Scientific$^{TM}$) was used as a molecular mass marker. This Figure extends **Fig. 3**.

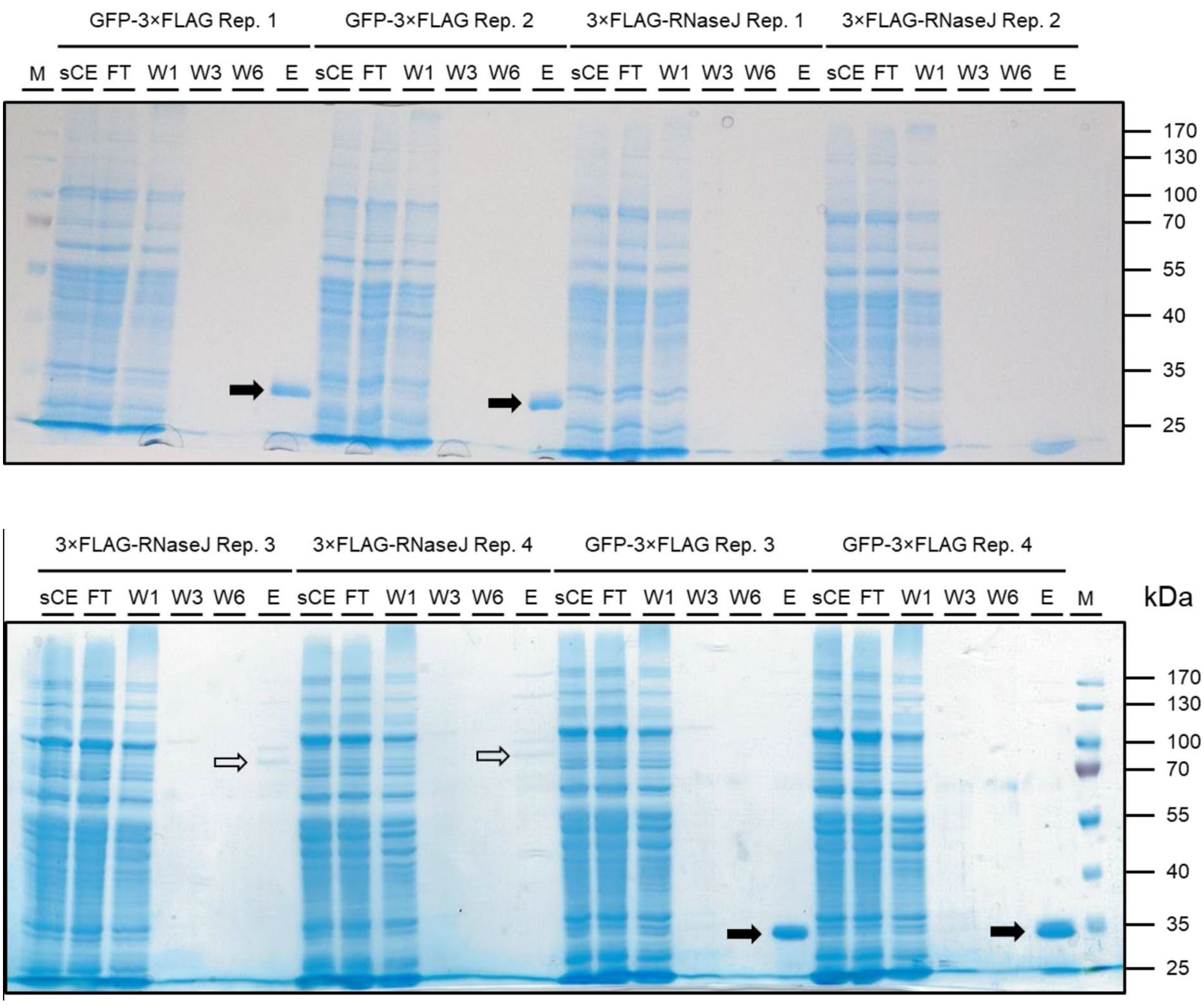


**Figure S2: Coomassie stained SDS-PAGE with samples from NTT RNase J co-immunoprecipitation.** *Synechocystis* strains carrying either pVZ322-Prha-*gfp*-*3×FLAG* or pVZ322-Prha-*3×FLAG*-*slr0551* were cultivated in the CellDeg system to obtain high density cultures. Four replicates were cultivated for each strain. Expression of RNase J was induced with 6 mg/mL L-rhamnose and harvested after 24 h. An equivalent of 1,000 OD750 units was lysed and FLAG-tagged proteins were purified with anti-FLAG M2 magnetic beads (Sigma Aldrich). Bound proteins were eluted with 0.4 µg/µL FLAG peptide. 2 µL of the soluble crude extract (sCE) and the flow through (FT) were loaded on a 10% SDS-PAA gel together with 16 µL of wash fractions (W1,3,6) and the elution (E). Proteins were stained with InstantBlue Coomassie Protein Stain (Abcam). M, PageRuler Prestained Protein Ladder. Bands corresponding to tagged GFP are marked by a black arrow, while empty arrows mark visible bands of tagged RNase J. This Figure extends data related to **Fig. 4**.

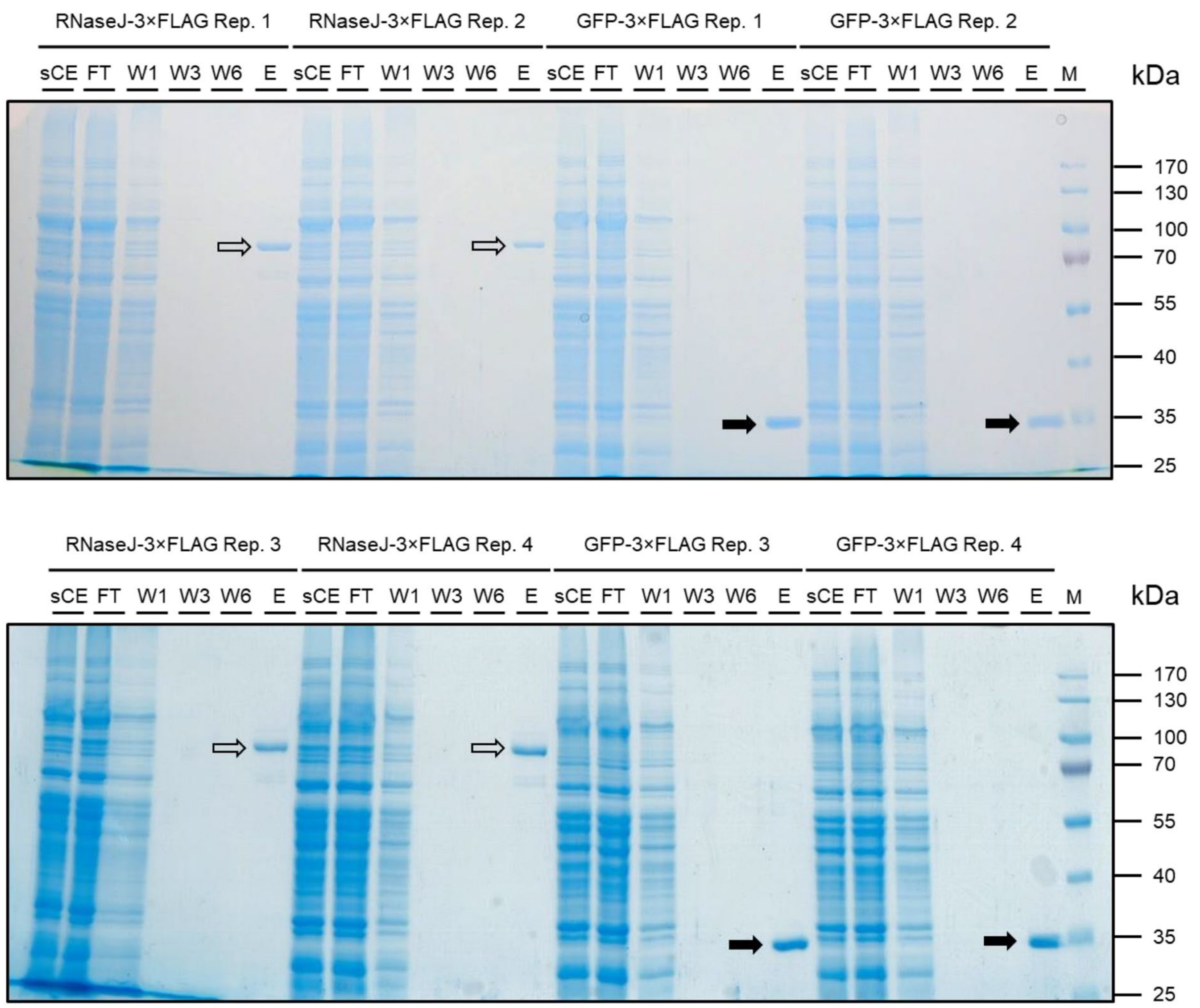


**Figure S3. Coomassie stained SDS-PAGE with samples from CTT RNase J co-immunoprecipitation.** *Synechocystis* strains carrying either pVZ322-P*petE*-*sfGFP*-*3×FLAG* or pVZ322-P*petE*-*slr0551*-*3×FLAG* were cultivated in the CellDeg system to obtain high density cultures. Four replicates were cultivated for each strain. Cells were induced with 0.5 mg/mL CuSO4 (2 mM) and harvested after 24 h. An equivalent of 1,000 OD750 units were lysed and FLAG tagged proteins were purified with anti-FLAG M2 magnetic beads (Sigma Aldrich). Bound proteins were eluted with 0.4 µg/µL FLAG peptide. 2 µL of the soluble crude extract (sCE) and the flow through (FT) were loaded on a 10% SDS-PAA gel together with 16 µL of wash fractions (W1,3,6) and the elution (E). Proteins were stained with InstantBlue Coomassie Protein Stain (Abcam). M, PageRuler Prestained Protein Ladder. Bands corresponding to tagged GFP are marked by a black arrow, while empty arrows mark visible bands of tagged RNase J. This Figure extends data related to **Fig. 4**.

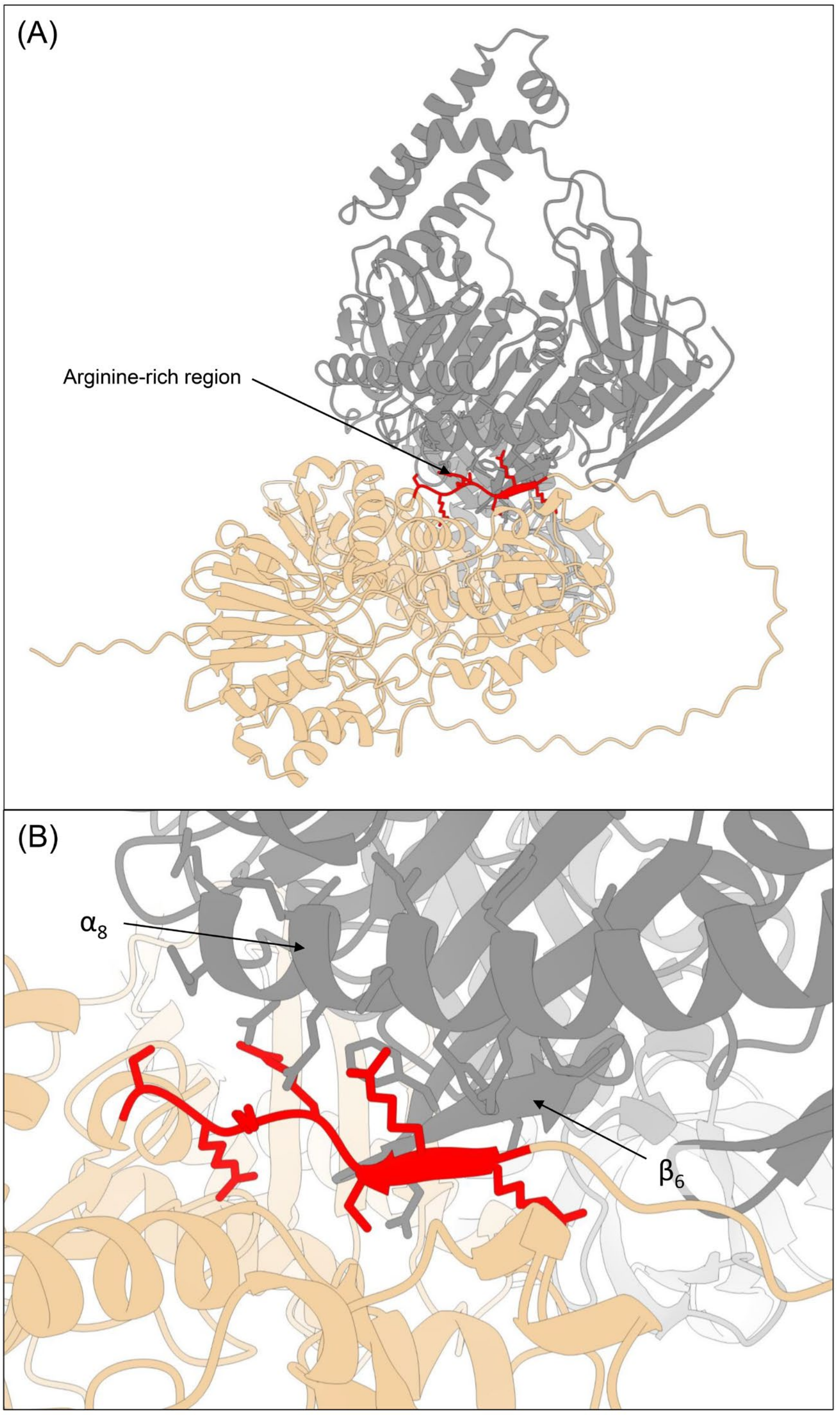


**Figure S4. Predicted interaction of RNase J and PNPase. (A)** Best-scoring structure of the RNase J-PNPase complex predicted by Alphafold 3 (Abramson *et al.* 2024). The pTM score was 0.52, the iPTM score 0.51. RNase J (light orange) has a small interaction surface with PNPase (gray). The arginine-rich motif at the CTD of RNase J is shown in red. **(B)** Enlargement of the interaction site. The arginine-rich motif of the CTD is in close proximity to a PNPase alpha helix ($\alpha_8$, residues 196–208) and a β-sheet ($\beta_6$, residues 158–161).

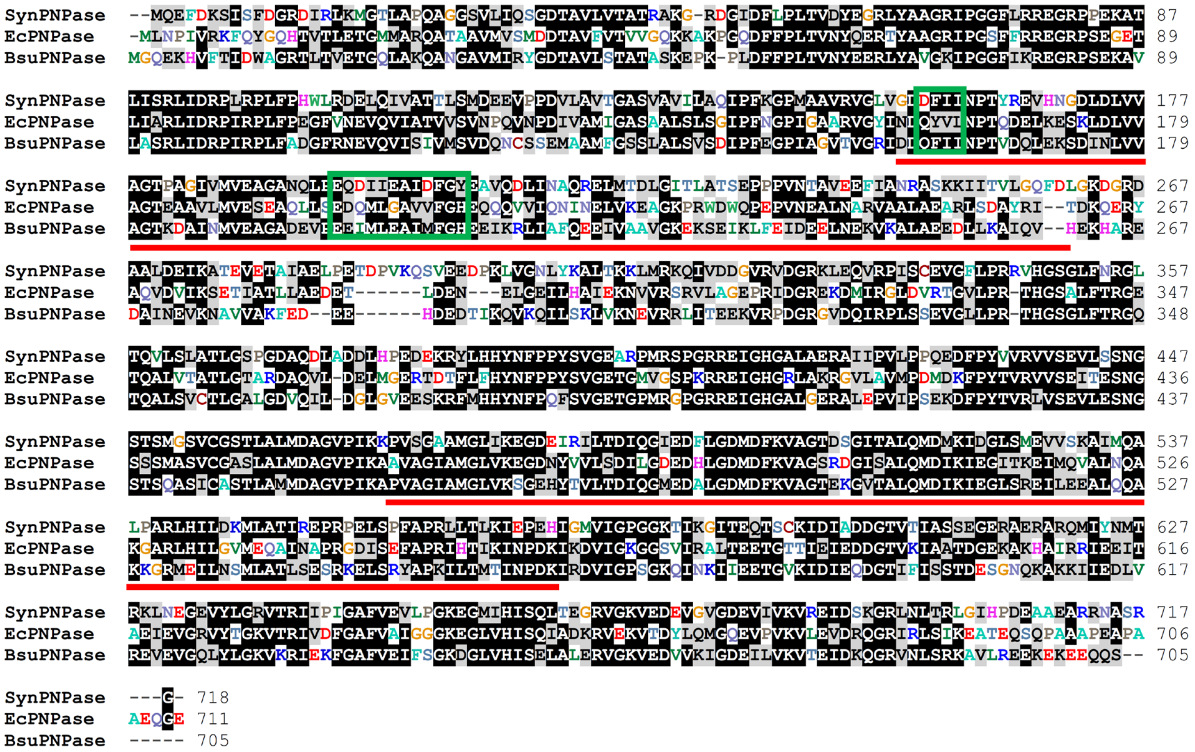

**Figure S5. Alignment of PNPase protein sequence**. The protein sequence of PNPase from *Synechocystis* sp. PCC6803 (Uniprot: P72659), *Escherichia coli* (B5YS54) and *Bacillus subtilis* (P50849) were aligned. Black shaded letters indicate conserved residues and grey shaded letters show similarity. Green boxes show residues which were in proximity to the arginine rich motif of the CTD of RNase J, observed in **Fig. S4**. Residues underlined in red show a previously identified interaction site which is important for the interaction of PNPase and RNase E in *E. coli* (Durán-Figueroa *et al.* 2006).

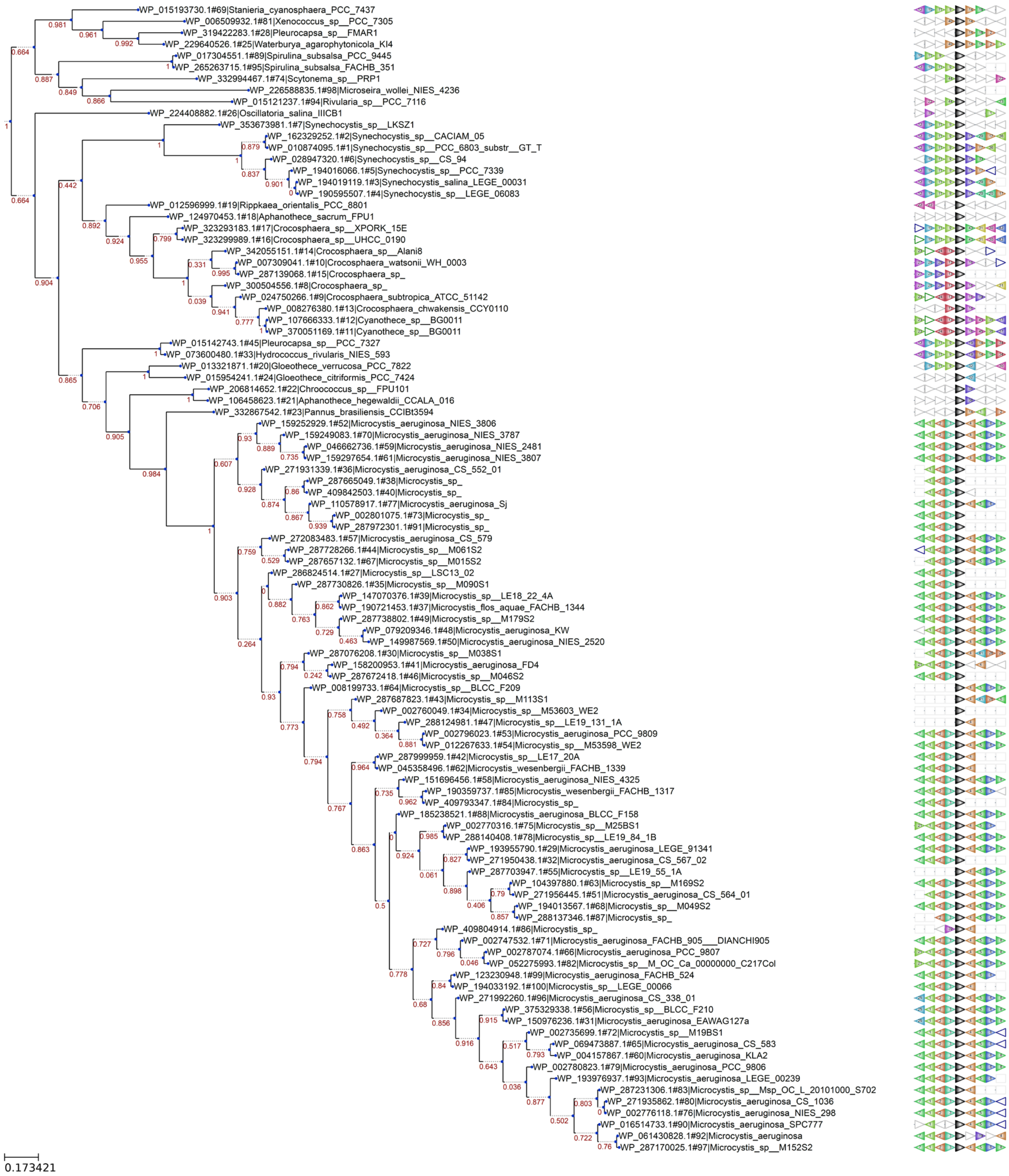


**Figure S6. Analysis of the genomic neighborhood of *slr0552*.** Synteny analysis of *slr0552* with webFlaGs (Saha *et al.* 2021) was performed to study conservation of the genomic neighborhood of *slr0552*. The refseq_protein (full) database was searched with a maximum output of 100 BlastP hits. Black arrowheads represent *slr0552* homologs. Light green arrowheads with the number 9 represent *rnj* homologs. 15 organisms (including *Synechocystis* 6803) have an *rnj* homolog upstream of *slr0552* or its respective homolog. Seven sequences belong to the genus *Synechocystis*. The same organization is also found in more distant cyanobacteria, including species from *Cyanosphaera*, *Spirulina*, *Crocosphaera*, *Pleurocapsa*, *Hydrococcus*, and *Gloeothece*.

### Supplementary Tables

**Table S1. List of oligonucleotides used in this work.**

| Oligo | Full Name | Sequence | Remark |
|---|---|---|---|
| RB 1 | petE promoter_fwd | ATCACGAGGCCCTTTCGTCTGAAGGGATAGCAAGCTAATTTTTATG | For reverse amplification of pUC19 with only essential elements for cloning. |
| RB 2 | petE promoter_rev | TTTTAGCCATACTTCTTGGCGATTGTATC | |
| RB 3 | RNAse J_fwd | GCCAAGAAGTATGGCTAAAAATACTCAAACC | Amplification of *slr0551* (*rnj*) sequence with 3´-end overhang to pUC19. |
| RB 4 | RNAse J_rev | ACGCCAGCAACGCGGCCTTTTTATCATCATCATCTTTATAATCAATATCATGATCTTTATAATCGCCATCATGATCTTTATAATCGGAAGAAACAGAAGTGGTAG | |
| RB 5 | Backbone pUC19_fwd | AAAGGCCGCGTTGCTGGC | For reverse amplification of pUC19 with only essential elements for cloning. |
| RB 6 | Backbone pUC19_rev | AGACGAAAGGGCCTCGTGATAC | |
| RB 7 | petE_RNAse J_3X FLAG tag_fwd | TATGCTCTTCTGCTCCTGCAGAAGGGATAGCAAGCTAATTTTTATG | For cloning of P*petE*-*slr0551*-*3×flag*-T*ooP* cassette into pVZ322. |
| RB 8 | petE_RNAse J_3X FLAG tag_rev | CTGCCCGGATTACAGATCCTAATAAAAAACGCCCGGCGGCAACCGAGCGAACTATTTATCATCATCATCTTTATAATCAATATC | |
| RB 9 | pet E RNase J ooP term_fwd | TGCTCCTGCAGAAGGGATAGCAAGCTAATTTTTATG | For cloning of P*petE*-*slr0551*-TooP cassette into pVZ322. |
| RB 10 | pet E RNase J ooP term_rev | CTGCCCGGATTACAGATCCTAATAAAAAACGCCCGGCGGCAACCGAGCGAACTAGGAAGAAACAGAAGTGGTAG | |
| RB 11 | FLAG/RNJ_fwd | AATATACAAAGGAGGTAGAAATGGATTATAAAGATCATGATGG | Amplification of 5´3×flag-*slr0551* with overhang to pUC19-Prham |
| RB 12 | FLAG/RNJ_rev | AATTTGGTACCGAGCTGCAGCTAGGAAGAAACAGAAGTG | |
| RB13 | pUC19_Rha_fwd | CTGCAGCTCGGTACCAAATTC | Amplification of pUC19-Prham vector |
| RB14 | Prha_slr0551_Frg1_rev | TATAATCCATTTCTACCTCCTTTGTATATTATAAACTTACC | |
| RB 15 | Rp_XmnI_Prha_fwd | GCCGCCCGCATTGGAGAAATAAGACCCCCGCACCGAAA | Amplification of P*rha*-flag-*slr0551*-Km-rhaS for cloning into XmnI digested pVZ322_short. |
| RB 16 | Rp_XmnI_Prha_rev | CAGATCGTTGACGAGTATTAAATAAAAAACGCCCGGCGGCAACCG | |
| RB17 | antiC3S1_fwd | CTTTAGGTGGGCGTTGACCT | Northern hybridization for CRISPR3 Spacer1-4 crRNA. From (Behler *et al.* 2018) |
| RB18 | antiC3S4_rev | TAATACGACTCACTATAGGGTAATAGTAATGACAGGCAG | |
| RB19 | 5S oligo 6803 | CTTGGCATCGGACTATTGTGCCGT | RNA quantification |
| RB20 | PrimerExt_Ladder_fwd | CTCGATGGCTTACTGTATAAG | DNA template for reference sequence ladder, primer extension. From (Behler *et al.* 2018) |
| RB21 | PrimerExt_S3_rev | CTACTAAGCTCGACAATTG | DNA template for reference sequence |

| | | | |
|---|---|---|---|
| | | | ladder, primer extension, annealing site in spacer 3. From (Behler *et al.* 2018) |
| RB22 | C3-Control-22nt | GUCUCCACUCGUAGGAGAAAUU | Synthetic ribonucleic acids for the preparation of the C3 ladder |
| RB23 | C3_24-30 deleted | GUCUCCACUCGUAGGAGAAAUUAGGAAAC | |
| RB24 | C3_5'-1 | UCUCCACUCGUAGGAGAAAUUAAUUGAUUGGAAAC | |
| RB25 | C3_unstructured | GGGAGAACACGUAGGAGAAAUUAAUUGAUUGGAAAC | |
| RB26 | C3_5'+A | AGUCUCCACUCGUAGGAGAAAUUAAUUGAUUGGAAAC | |
| RB27 | C3-spacer2-repeat | AGCGCCACAGCUGACAGAGUUCCUGAAGGAAGCUAAGUCUCCACUCGUAGGAGAAAUUAAUUGAUUGGAAAC | |

**Table S2. Mass spectrometry results of co-immunoprecipitation with N- and C-terminally tagged RNase J.**

**See separate Excel file**

**Table S3.** Protein class attribution of RNase J-coenriched proteins. The 20 and 36 proteins enriched in the co-IP with NTT, or CTT RNase J, respectively, were analyzed with PANTHER (Thomas *et al.* 2022) and classified according to their protein class category. Protein classes are shown in the middle. Percentage of protein class attributions identified in the NTT RNase J co-IP are shown in the left column, and those identified in the CTT co-IP in the right column.

| NTT RNase J | Protein class | CTT RNase J |
|---|---|---|
| 10.00% | chaperone (PC00072) | 5.90% |
| 0.00% | DNA metabolism protein (PC00009) | 11.80% |
| 5.00% | gene-specific transcriptional regulator (PC00264) | 8.80% |
| 35.00% | metabolite interconversion enzyme (PC00262) | 14.70% |
| 5.00% | protein-binding activity modulator (PC00095) | 2.90% |
| 0.00% | protein modifying enzyme (PC00260) | 8.80% |
| 5.00% | RNA metabolism protein (PC00031) | 2.90% |
| 0.00% | translational protein (PC00263) | 5.90% |
| 5.00% | transporter (PC00227) | 2.90% |
| 35.00% | unclassified | 35.30% |

### Supplementary Methods

**In vitro cleavage assay.** 5 pmol of synthetic oligoribonucleotide RB24 (C3 spacer 2-repeat or RB23 (C3 repeat + 5´A) were incubated with approximately 400 ng of recombinant RNase J and/ or RNase E in different combinations in RNase E cleavage buffer (Behler *et al.* 2018) in a 10 µL reaction volume. Samples were incubated at 30°C for 30 min. Reaction was stopped by adding 10 µL of 2 × RNA loading dye and incubated at 70°C for 10 min for denaturation. Samples were then loaded on an 8M urea - 15% PAA gel. Gel was stained with SYBR Gold (Thermo Fisher Scientific) at a 1:10,000 dilution with 0.5× Tris–borate–EDTA buffer. Signals were detected with a Laser Scanner Typhoon FLA 9500 (GE Healthcare) with the following settings: excitation, 473 nm; emission filter long pass blue ≥ 510 nm; photomultiplier value, 600.

**Preparation of alkaline hydrolysis ladder.** 10 pmol $\mu L^{-1}$ of synthetic oligoribonucleotide (C3 spacer2-repeat or C3 repeat + 5´A) were incubated with 50 mM $NaHCO_3$, pH 9.3. The reaction was incubated for 5 min at 95°C, placed on ice for 1 minute and RNA loading dye was added to the mix.

**Preparation of the C3 ladder.** The synthetic RNAs RB22-RB27 were mixed as follows: RB22, 1.41 µL; RB23, 1.07 µL; RB24, 0.89 µL; RB25, 0.86 µL; RB26, 0.84 µL and RB27, 0.43 µL. The RNA mixture was added to 14.4 µL of RNase-free water and 20 µL of 2× RNA loading dye.
